# A paradigm for post-embryonic Oct4 re-expression: E7-induced hydroxymethylation regulates Oct4 expression in cervical cancer

**DOI:** 10.1101/2023.04.10.536212

**Authors:** Theofano Panayiotou, Marios Eftychiou, Eleutherios Patera, Vasilis Promponas, Katerina Strati

## Abstract

The Octamer-binding transcription factor-4 (Oct4) is upregulated in different malignancies, yet a paradigm for mechanisms of Oct4 post-embryonic re-expression is inadequately understood. In cervical cancer, Oct4 expression is higher in HPV-related than HPV-unrelated cervical cancers and this upregulation correlates with the expression of the E7 oncogene. We have reported that E7 affects the Oct4-transcriptional output and Oct4-related phenotypes in cervical cancer, however, the underlying mechanism remains elusive. Here, we characterize the Oct4-protein interactions in cervical cancer cells and reveal that Methyl-binding proteins (MBD2 and MBD3), are determinants of Oct4 driven transcription. E7 triggers MBD2 downregulation and TET1 upregulation, thereby disrupting the methylation status of the Oct4 gene. This coincides with an increase in the total DNA hydroxymethylation leading to the re-expression of Oct4 in cervical cancer and likely affecting broader transcriptional patterns. Our findings reveal a previously unreported mechanism by which the E7 oncogene can regulate transcription by increasing DNA hydroxymethylation and lowering the barrier to cellular plasticity during carcinogenesis.

**Teaser:** E7 modulates Oct4 interactions and related characteristics in cervical cancer cells by altering the DNA methylome.

## Introduction

The stem cell-related transcription factor, Octamer binding transcription factor-4 (Oct4), with well-established traits during embryogenesis and pluripotency maintenance [1], has been shown to be expressed in certain malignancies (hepatocellular [2], cervical [3], head and neck [4], ovarian [5], breast [6] and prostate cancer [7]). We have previously shown that Oct4 is more highly expressed in Human Papillomavirus (HPV)-dependent compared to HPV-independent cervical cancer cells and this upregulation correlates with the expression of the HPV E7 viral oncogene [8]. Proteomic and genomic approaches have provided insights on the role of Oct4 in pluripotency and stemness, nevertheless these approaches remain a challenge for elucidating a role in cancer due to lower expression levels of Oct4 in cancer cells and tissues. In embryonic stem cells (ESC), Oct4 expression is tightly regulated by signaling molecules and epigenetic regulators to retain the balance between pluripotency and differentiation [9]. Oct4 expression and function vary across cell types [10–14] and depend on the chromatin state, which determines the balance between pluripotency and differentiation in ESCs. ESCs remain pluripotent when the Oct4 locus is hypomethylated whereas hypermethylation of Oct4 leads to the differentiation of ESCs [15]. Other key molecules that alter the chromatin structure and Oct4 expression are epigenetic regulators. An important regulator complex is the Nucleosome Remodeling and Deacetylase complex (NuRD), which alters the epigenetic definition of Oct4, mediating transcriptional repression in ESCs during differentiation [16, 17]. Additional evidence shows that not only the NuRD complex can regulate the expression profile of Oct4 in ESCs but also Oct4 recruits the NuRD complex on regulatory regions of Oct4-target genes to control their status (active/inactive) [18, 19]. The repressive NuRD complex is required for many physiological functions apart from regulating pluripotency [20] such as the regulation of B-and T-cell development [21], and the differentiation of cells with hematopoietic lineages [22]. Despite its involvement in normal developmental processes, the NuRD complex is associated with the onset and progression of cancer, with many of its components aberrantly expressed in cancer tissues [23]. In cervical cancer, there is evidence that the main HPV oncogene, the E7, interacts with NuRD components such as Histone deacetylase 1/2 (Hdac1/2) via the Chromodomain helicase DNA binding protein 4 (Chd4) to provide a proliferative advantage by driving cells into the S-phase of the cell cycle [24] while Proline Rich methyltransferase 5 (PRMT5) associates with the NuRD complex to increase the invasive potential and maintain DNA methylation in cervical carcinoma [25].

Epigenetic DNA alterations are recognized as one of the hallmarks of cancer development. In cervical cancer, these epigenetic modifications are mainly driven by DNA methylation or demethylation, chromatin remodeling and histone modifications. High levels of DNA methylation are commonly observed in most cervical tumors while the methylation status of cervical lesions is considered as a marker for this cancer [26]. Upon HPV infection, the viral and host DNA methylation facilitates viral persistence and carcinogenesis. The grade of HPV L1 methylation is indicative of disease severity and persistent HPV infection in the cervix [27], while persistent HPV infection is due the hypermethylation of the E2 binding site on the Long Control region (LCR) [28]. E6 and E7 known for binding and degrading p53 and pRb respectively, activate DNA-methyltransferases like DNMT1 to trigger hypermethylation and promote the repression of tumor-suppressor genes of the host [29]. Apart from viral and host DNA methylation, demethylation might sometimes contribute to the onset and progression of cervical cancer. For instance, in W12-immortalised keratinocytes the binding site of E2 on the LCR is demethylated leading to the continuous activation of viral oncogenes [30, 31].

Apart from cervical tumors, aberrant DNA methylation is evidenced in many cancer types and is correlated with transcriptional repression [32], although hydroxymethylation, which is the first derivative of the active demethylation cycle and is known to be associated with active transcription [33], has gained increasing attention as a biomarker in premalignant or metastatic lesions in different cancer types [34–36]. For instance, in hepatocellular carcinoma, the oxidized 5-hydroxymethycytosine (5hmC) levels are upregulated in premalignant lesions while during advanced malignancy 5hmC levels fade away and are substituted by high levels in 5-methylcytocine (5mC) [37]. On the contrary, metastatic prostate tissues expressed higher total 5hmC compared to the normal or pre-cancerous state [38]. The driving force behind the elevated hydroxymethylation in either premalignant or metastatic lesions is the activation or re-expression of the TET-eleven translocation enzymes (TET), which carry out the oxidation of 5mC to 5hmC [35, 39]. On the chromatin assembly, hydroxymethylation is strongly associated with enhancers and gene bodies whereas methylation is mostly seen on gene promoters. In ESC the MBD2 and MBD3 components of the NuRD are highly associated with hypermethylated or hydroxymethylated loci respectively, affecting the activation or inactivation of certain genes.

Here, we describe the underlying mechanism by which Oct4 is re-expressed in cervical cancer by identifying the Oct4-protein-protein interaction (PPI) network in C33A cervical cancer cells. We established the Oct4-NuRD interaction in cervical cancer while we have determined that the E7 oncogene alters the binding from MBD2 to MBD3 on the Oct4-NuRD complex. Additionally, we provide evidence that E7 affects the binding and regulation of MBD2 on the Oct4 gene maintaining cell viability. We thus propose a unique contribution of the viral oncogene E7 in regulating global DNA (de)methylation dynamics in cervical cancer cells, and a potential paradigm of Oct4 re-expression in post-embryonic cells.

## Results

### Oct4 interacts with the NuRD complex in cervical cancer

Currently evidence regarding Oct4 protein-protein interactions derives from studies in ESC and Induced Pluripotent Stem cells (iPSC) [19, 40, 41] however, the Oct4 interaction network in cancer remains uncharacterized. To profile the protein-protein interactions of Oct4 in C33A cervical cancer cells (HPV-negative) we performed antibody-mediated Affinity Purification coupled to Mass Spectrometry (AP-MS). To enhance the capacity for antibody mediated-purification, we transiently overexpressed Oct4 in cervical cancer cells (table S1) For negative controls, we used an IgG antibody for immunoprecipitation and the Oct4-knockdown C33A transduced cells [8]. For the analysis, we used a cut-off p-value less than 0.05 (p<0.05) and a fold change greater than 2 (FC>2). As anticipated, Oct4 (POU5F1) was found significantly enriched in C33A transfected cells while we identified 1605 enriched hits as indicated by the volcano plot in Figure 1A. We validated four enriched hits, MCM7, PCNA, Vimentin and p53 via immunoprecipitation followed by Western Blot (Fig. 1B). In agreement with the Mass-spectrometry findings, Oct4 immunoprecipitated with these four proteins while interaction was absent in the shOct4-knockdown cells and the IgG negative controls.

**Fig. 1.**
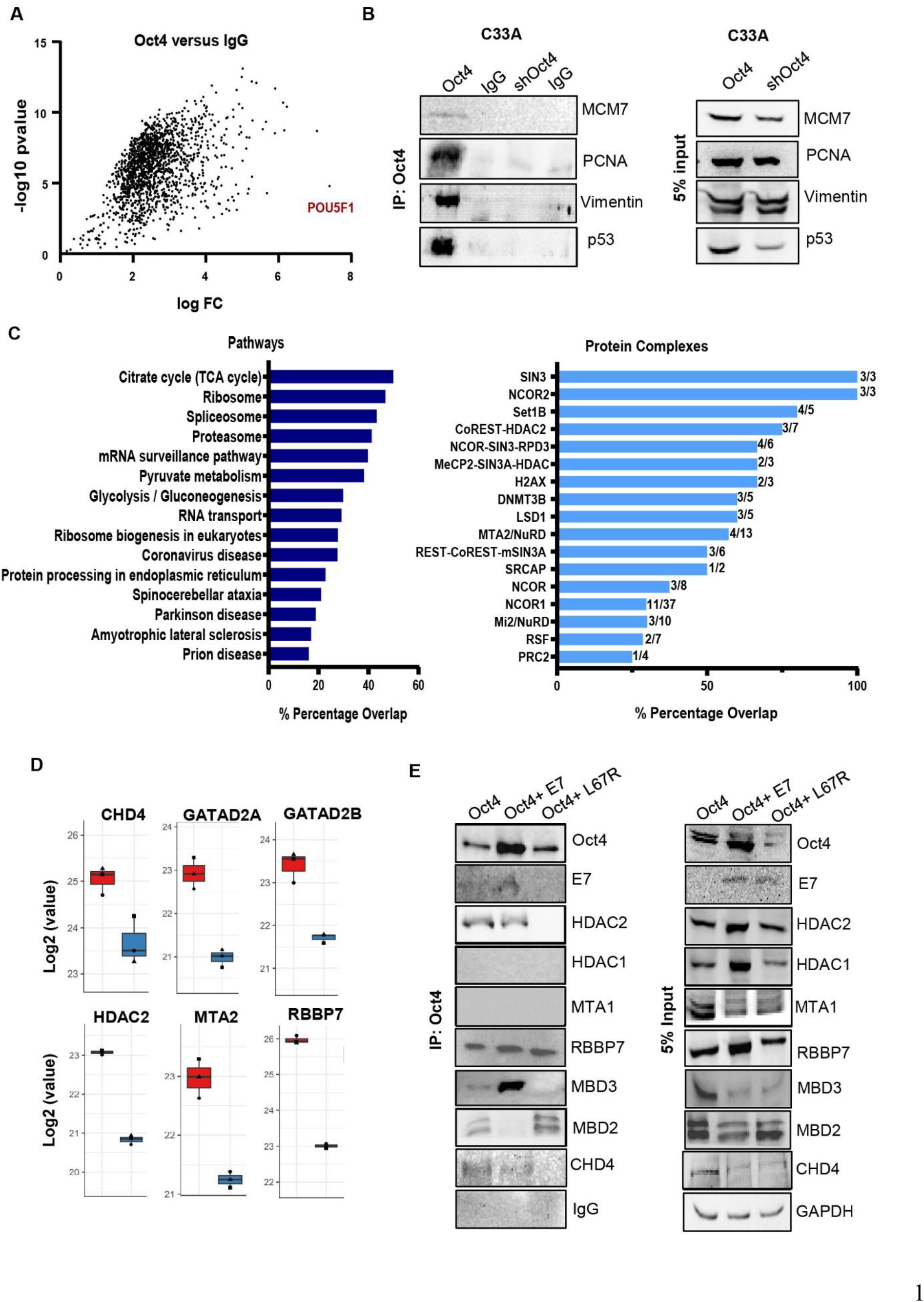
The Oct4 interaction with MBD proteins in cervical cancer is regulated by E7. (A) Volcano plot indicating 1605 proteins interacting with Oct4 in C33A cervical cancer cells as revealed by Mass Spectrometry. IgG was used as the negative control of the experiment, and three independent replicates were used. (B) Validation of Mass spectrometry through the immunoprecipitation of Oct4 in C33A and C33A Oct4-knockdown cells followed by western blot. IgG was used as the negative control and 5% input was used to verify the expression of these proteins in the cells. (C) Pathway analysis performed by the Enrichr Software demonstrates the pathways regulated by Oct4 interactors (dark blue color) while protein complexes involved in epigenetic regulation, which interact with Oct4, are shown in light blue color. (D) Components of the NuRD complex interact with Oct4 as shown by Mass spectrometry while the red bars indicate Oct4 immunoprecipitation and blue bars indicated IgG immunoprecipitation. (E) C33A cells transfected with Oct4, Oct4 & HPV16 E7 and Oct4 & E7 L67R vectors were used for the Immunoprecipitation experiment. Oct4 was pulled down with an Oct4 antibody and Western blot was performed to reveal Oct4 interactions with NuRD-associated proteins. IgG was used as the negative control of the experiment while 5% input verifies the expression of these proteins in cells.

To bolster our proteomic approach, we performed computational analyses on publicly available PPI datasets. We generated an Oct4-PPI network by mining and processing data from existing literature using 3 different databases (APID, IntAct and BioGRID [42–44]). 554 proteins were found to interact with Oct4, while complementary to the Oct4-PPI analysis we generated an E7-PPI network to identify putative common interactors of both Oct4 and the HPV oncogene E7. Further computational analyses revealed 41 proteins as reported interactors between both Oct4 and E7 (fig S1A). To examine the molecular and functional interplay between the Oct4-E7-PPI network, enrichment analysis was achieved. Notably, Oct4 and E7 shared PPIs exist in several complexes, the majority involved in epigenetic regulation, histone modifiers or histone re-modelers, as molecules that regulate the transcriptional process. The NuRD complex emerges from this analysis as the one containing most common interactors between Oct4 and E7 (fig S1B). Even though this complex is associated with developmental processes, recently it has been implicated in the process of carcinogenesis [45–48].

To investigate which pathways may be regulated by Oct4 based on its protein-protein interactions in C33A cells, we performed pathway analysis using the Enrichr software (Fig. 1C). Pathways predicted to be regulated by Oct4-PPIs include metabolic pathways and pathways implicated in protein synthesis and processing. Additionally, we examined whether Oct4 identified interactors participate in protein complexes in cervical cancer cells and cross-referenced these complexes with our computational analyses. Interestingly, we detected the NuRD complex to be also regulated by Oct4 in cervical cancer cells (Fig. 1C) while many components of the NuRD complex immunoprecipitated with Oct4 such as CHD4, GATAD2A, GATAD2B, HDAC2, MTA2 and RBBP7 (Fig. 1D). We have previously reported an interaction between Oct4 and the viral oncogene E7 in cervical cancer cells. This interaction was not detected when the aminoacid 67 was mutated (L67R E7 mutant) [8]. To validate the Oct4-NuRD interaction in C33A cells and further examine whether this interaction is affected by the presence of the viral oncogene E7, we transfected C33A cells with Oct4, Oct4+E7 and Oct4+L67R vectors and following Oct4 immunoprecipitation, western blot was carried out to validate the Oct4-NuRD interaction. Remarkably, we found that most interactions of Oct4 with components of the NuRD complex do not vary in the presence of E7. However, we found the readers of the complex known for their Methylated DNA binding domain (MBD), to vary in the presence of wildtype E7. MBD2 interacts with Oct4 only when the wildtype E7 is not expressed in C33A cells whereas upon E7 expression MBD2 no longer immunoprecipitates with Oct4. Instead, in the presence of E7, MBD3 immunoprecipitates with Oct4 (Fig. 1E). From previous studies we know that MBD2 and MBD3 are mutually exclusive members of the NuRD complex [49], validating our co-immunoprecipitation data. Additionally, both reader MBD proteins bind genomic loci based on their methylation status [48, 50–53]. Hence, to investigate whether the E7 oncogene has an impact on pathways related to DNA methylation/demethylation in C33A cells, we isolated RNA from Oct4-expressing and Oct4+E7-expressing C33A cells for Quant sequencing. We used a cut-off p-value less than 0.05 (p<0.05) and we have identified 1134 differentially expressed genes (453 upregulated and 681 downregulated) (fig S1C). Specific highly upregulated and highly downregulated genes were selected for validating the sequencing by performing qRT-PCR (fig S1D) whereas RT-PCR was performed to validate the successful transfection of E7 (fig S1E). Pathway analysis illustrated several biological processes regulated by Oct4 and E7 such as metabolic processes, cancer and RNA surveillance and transport (fig S1F). Notably, we identified the mechanism of Base excision repair (BER) to be controlled by both Oct4 and E7. The BER mechanism is involved in the dynamic deletion of epigenetic marks on chromatin structure converting methylated cytosines into their unmethylated state. This conversion is part of the active demethylation process in the cells which transform 5-methylcytosine (5mC) to 5-hyroxymethylcytosine (5hmC), 5-formylcytosine (5fC) and 5-carboxylcytosine (5caC). 5fC and 5caC are converted to unmethylated cytosine through this BER system [54, 55].

### E7 elevates TET1 expression and global hydroxymethylation in C33A and HaCaT cells

E7 modulates epigenetic marks on chromatin via enzymes involved in the process of methylation and acetylation to regulate transcription. Here, we propose a new function of E7 in modifying methylation dynamics in C33A cells through the involvement of TET enzymes which catalyse the oxidation of 5mC to 5hmC, to 5fC and finally to 5caC (fig S2A).

Cervical cancer cells (C33A) and Human Immortalised keratinocytes (HaCaT) transfected with an Empty control (Neo Bam), HPV16 E7 and the mutant L67R were collected for assessing TET1 mRNA and protein levels. C33A and HaCaT cells expressing the wildtype but not the mutant E7 show an elevated expression of TET1 both at the mRNA and protein level. (Fig. 2, A-C and 2, F-H). Since E7 increases TET1 we investigated the possibility of an increase in the total 5-hydroxymethylcytosine levels (5hmC). Thus, we conducted a dot blot analysis by using C33A cells transfected with the Neo Bam Empty, HPV16 E7 and E7 L67R vectors. Extracted Genomic DNA was applied at two different concentrations (100ng and 10ng) on a nitrocellulose membrane in the form of a dot. At 100ng we detected an increase in 5hmC expression both in the presence of wildtype and mutant E7 compared to the empty control. The relative 5hmC levels in C33A were quantified and plotted (Fig. 2D). Additionally, we used immunofluorescence to validate the 5hmC profile in C33A upon the presence of E7. Both 5hmC and 5mC antibodies were used to stain the cells since an increase in 5hmC reflects a decrease in 5mC and vice versa. We observed a slight increased 5hmC intensity in HPV16 E7-C33A and E7 L67R-C33A cells compared to the Empty-C33A cells, even though no major difference in 5mC intensity was reported (Fig. 2E).

**Fig. 2.**
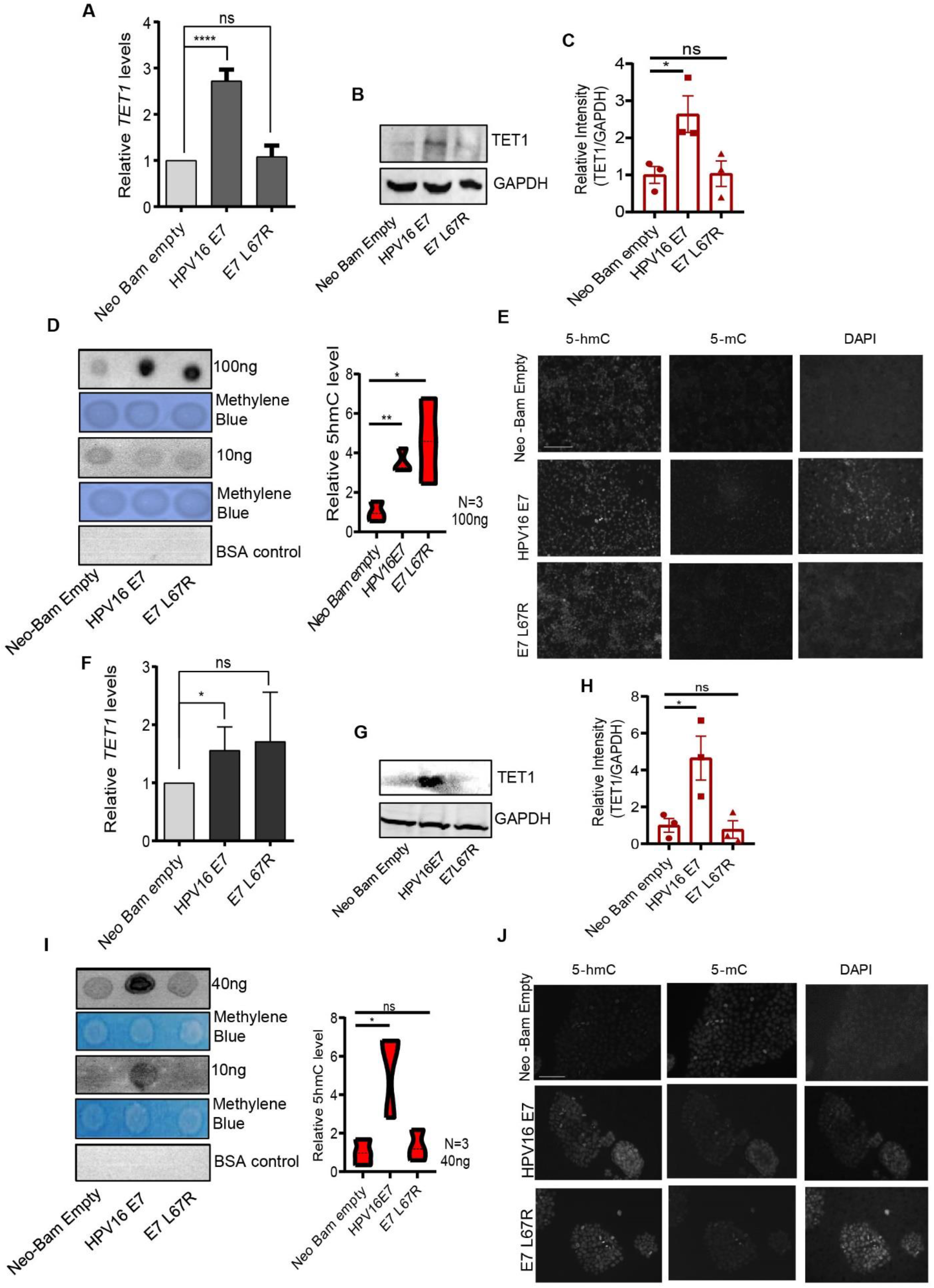
E7 increases TET1 expression and global hydroxymethylation in C33A and HaCaT cells. C33A cells transfected with Neo Bam empty, HPV16 E7 and E7 L67R plasmids were used to check TET1 (A) mRNA and (B) protein levels. GAPDH was used as a control. (C) Bar chart illustrating the average relative protein expression of TET1 compared to GAPDH. (D) Dot blot experiment illustrates 5hmC levels in C33A cells transfected with Neo Bam empty, HPV16 E7 and E7 L67R vectors. Genomic DNA was used at concentrations 100ng and 10ng. Methylene blue was used as a loading control whereas BSA was used as a negative control of the experiment. The violin plot demonstrates the relative 5hmc levels compared to methylene blue stain. (E) Immunofluorescence analysis shows 5hmC and 5mC expression in C33A cells, while DAPI was used to stain cell nuclei (Scale bars: 50μm). HaCaT transfected with Neo Bam empty, HPV16 E7 and E7 L67R vectors were used. TET1 (F) mRNA and (G) protein levels were investigated. GAPDH was used as a control. (H) Bar chart illustrating the average relative protein expression of TET1 compared to GAPDH. (I) Dot Blot experiment revealed 5hmC levels in HaCaT cells transfected with Neo Bam empty, HPV16 E7 and E7 L67R vectors. Genomic DNA was used at two concentrations (40ng and 10ng.) Bar chart demonstrates the relative 5hmc levels compared to methylene blue stain. (J) Immunofluorescence shows 5hmC and 5mC expression in HaCaT cells. Three independent replicates were used and plotted values on graphs are the Mean±SEM. The statistical analysis was performed with two-tailed Unpaired T-test with p<0.005 (ns = non-significant, *p<0.05, **p<0.01, ***p<0.001, ****p<0.0001).

Similarly, we used HaCaT cells transfected with Neo Bam Empty, HPV16 E7 and E7 L67R vectors for dot blot analysis revealing an increase in 5hmC levels upon the presence of wildtype E7 at 10 and 40ng (Fig. 2F). Immunofluorescence analysis validated the global increase in 5hmC and decrease in 5mC in E7-expressing HaCaT cells compared to the Empty control and the mutant E7 (Fig. 2J). To further confirm the impact of E7 in the demethylation cycle, we analysed the mRNA expression of certain genes participating in the demethylation process as revealed by the Quant-seq data. Upon the presence of E7, we noticed an upregulation of the *TDG* enzyme (thymine DNA glycolase) and its interactor *GADD45A* [56] which participate in the active demethylation cycle and take part in the BER pathway (fig S2A). Apart from DEGs illustrated by the Quant-seq, we checked the profile of certain enzymes known for their methylating activities observing a significant downregulation in DNA methyltransferase enzymes (*DNMT1*, *DNMT3A* and *DNMT3B*) (fig S2B). Thus, we propose the contribution of the viral oncogene E7 in the active demethylation cycle (fig S2A).

### E7 alters the methylation status of the human Oct4 gene

To address the impact of E7 on hydroxymethylation locally, we conducted DNA-methylation and DNA-hydroxymethylation Chromatin immunoprecipitation on the human Oct4 gene. For this reason, we transfected C33A cervical cancer cells with the Neo Bam Empty and HPV16 E7 vectors while cells were fixed and collected for Chromatin shearing and for verifying successful transfection (fig S3A, B). We observed a significant increase in the methylation state of 10 randomly selected loci on the hOct4 gene in Neo Bam Empty-expressing C33A cells compared to the IgG control (Fig. 3A). These ten loci are spanning various regions on the hOct4 gene (including upstream and downstream the Transcription Start Site TSS) (fig S3C). E7-expressing C33A cells displayed no significant change in the methylation levels of the ten loci tested when compared to the IgG control (Fig. 3A).

**Fig. 3.**
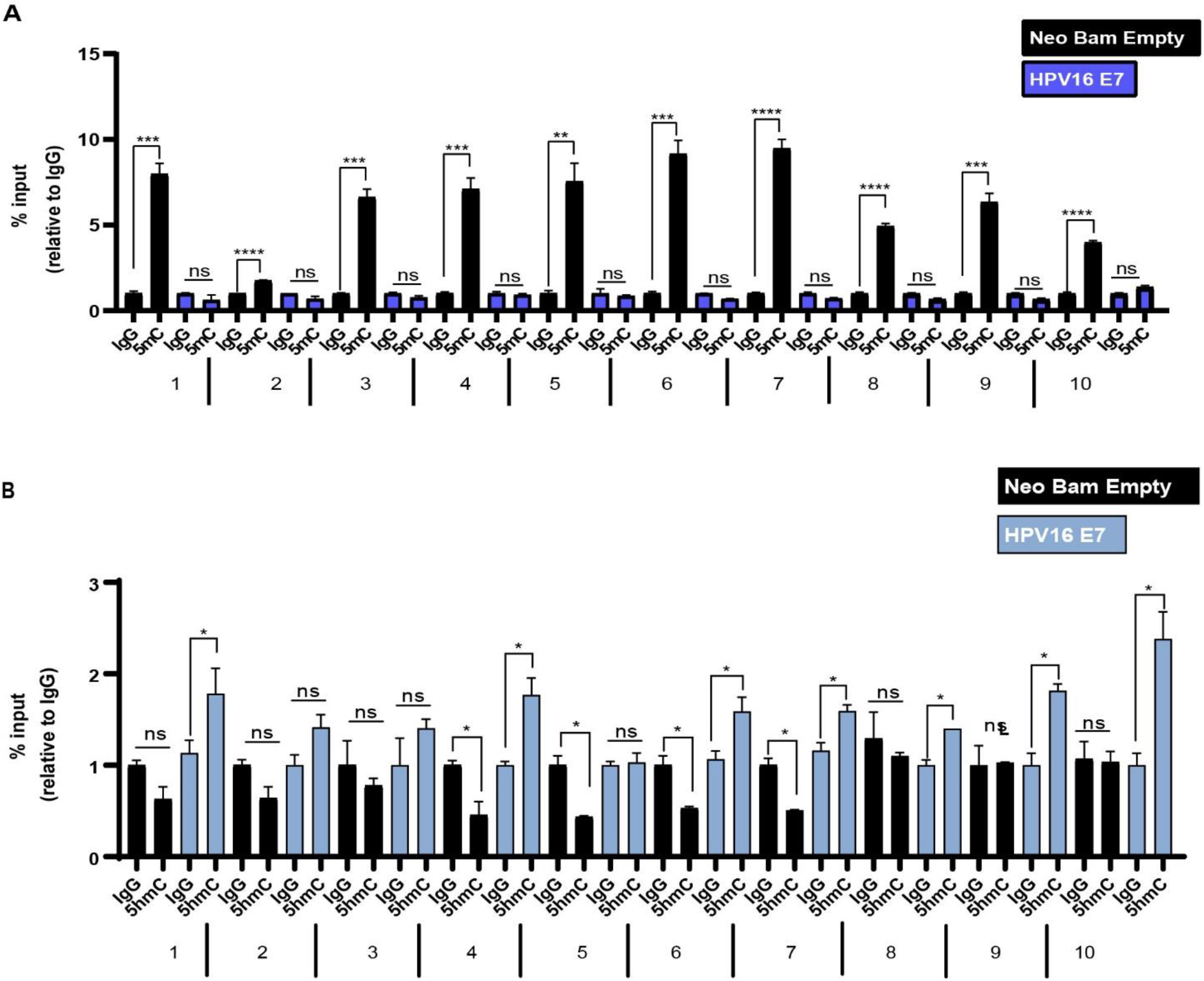
E7 expression in C33A cells deregulates the methylation status on the human Oct4 gene. C33A cells transfected with Neo Bam empty and HPV16 E7 vectors were fixed, and DNA was sheared for (A) methylation DNA immunoprecipitation and (B) hydroxymethylation DNA immunoprecipitation. IgG was used as the negative control of the experiment. qRT-PCR was performed to check 5mC and 5hmC enrichment by using primers targeting different loci on the hOct4 gene. Three independent replicates were used and plotted values on graphs are the Mean±SEM. The statistical analysis was performed with two-tailed Unpaired T-test with p<0.005 (ns = non-significant, *p<0.05, **p<0.01, ***p<0.001, ****p<0.0001).

In Hydroxymethylation DNA immunoprecipitation experiments, we found an enrichment on certain loci on the hOct4 gene only in E7-expressing C33A cells. These enriched loci are found closer to TSS and downstream the TSS (very few loci were enriched upstream the TSS). (Fig 3B).

### MBD2 downregulation modifies global hydroxymethylation in cervical cancer cells

MBD proteins are known for binding methyl regions of chromatin, nonetheless, one MBD family member, the MBD3, cannot make a strong contact with methylated chromatin regions, instead it binds tightly unmethylated or hydroxymethylated loci [53, 57]. For this reason, we assessed MBD3 as a reader of E7-induced hydroxymethylation in cells. We transfected both HaCaT and C33A cells with Neo Bam Empty, HPV16 E7 and E7 L6R plasmids while cells were assessed for MBD3 mRNA and protein expression. Surprisingly, we noticed that MBD3 mRNA expression was not affected by the presence of either the wildtype of the mutant E7 in C33A cancer cells but not in keratinocytes (Fig. 4A). This finding led us hypothesize that the elevated 5hmC expression in E7-expressing cells could be due to a decrease in their methylation status. Consequently, we checked the expression profile of MBD2 upon the presence of E7. Notably we observed a significant downregulation of MBD2 mRNA and protein levels in E7-expressing HaCaT and C33A cells (fig 4b and sup fig 4A. The reduced MBD2 pattern detected in the L67R-expressing C33A cells (Fig. 4B) possibly explains the increase in global 5hmC expression in L67R-transfected C33A cells (Fig. 2D). To further ensure that the reduced MBD2 expression led to an increase in global hydroxymethylation, we performed a dot blot analysis in two cervical cancer cells lines (C33A and CaSki) transduced with shRNA MBD2 knockdown construct and the shluciferase control. While validating the MBD2 knockdown at the mRNA and protein level, we noticed that shMBD2-2 in C33A cells could not yield a reduced expression of MBD2 (fig. S4A, B). For this reason, we chose to omit these cells from further analyses. We observed that shMBD2-1-transduced C33A cells displayed an increased 5hmC expression compared to the shluciferase-transduced C33A cells. Similarly, in shMBD2-knockdown CaSki cells, there was an increase in global 5hmC profile compared to the shluciferase control both at the 25ng and 100ng that we have tested (Fig. 4E) These differences in 5hmC levels were quantified and plotted in the form of a violin plot (Fig. 4, C and E). Lastly, we performed an immunofluorescence analysis to verify the elevated expression of 5hmC observed by dot blots by using shMBD2-knockdown C33A and CaSki cells. Again, we found a slight increase in the intensity of 5hmC in shMBD2 knockdown cells compared to the shluciferase control (Fig. 4, D and F). While quantifying the changes in intensity observed in C33A and CaSki cells in respect to 5hmC we noticed a higher intensity in the shMBD2 knockdown cells compared to the control, however the statistical trend was insignificant (fig S4C).

**Fig. 4.**
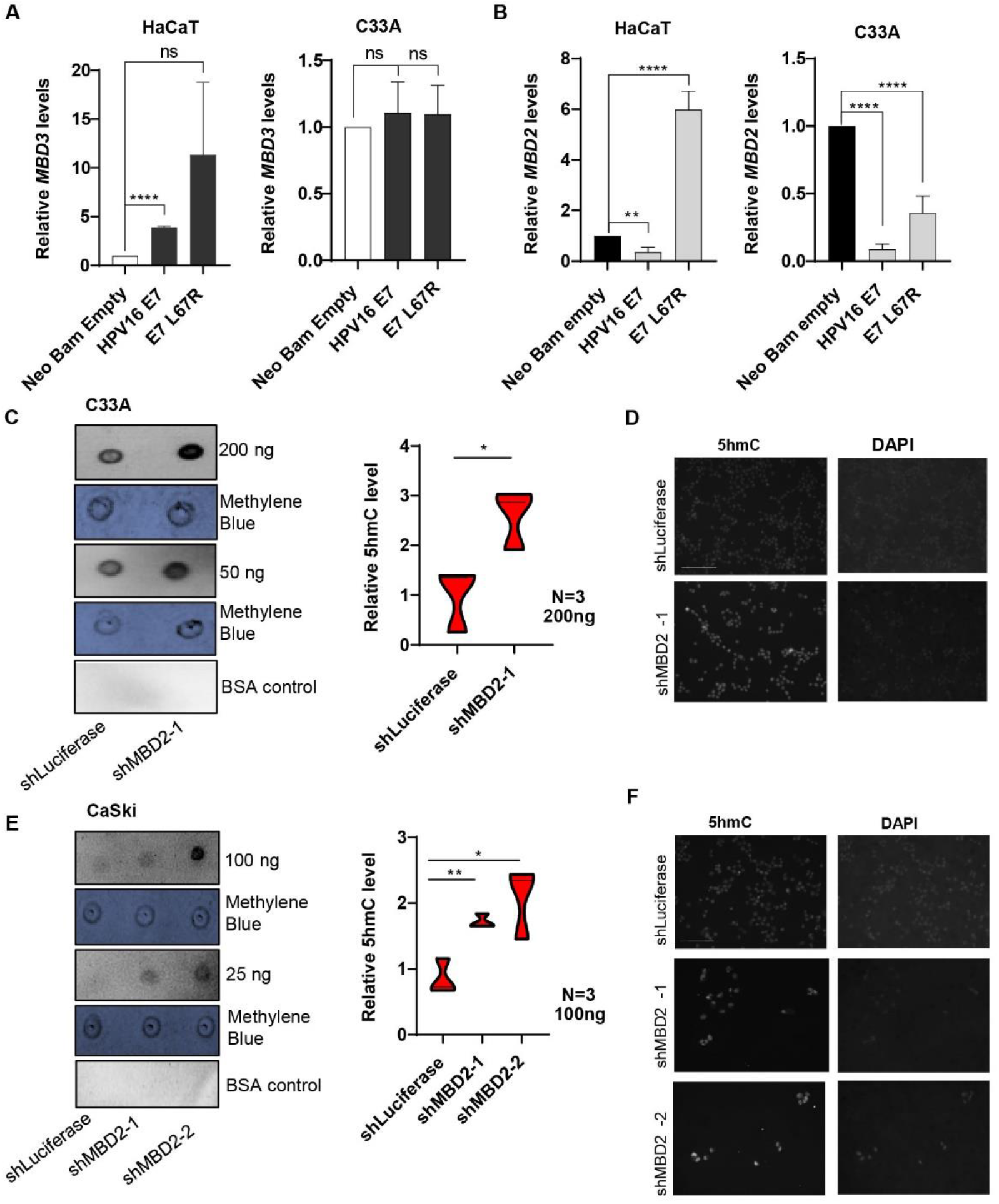
MBD2 knockdown increases global 5hmC levels in cervical cancer cells. HaCaT and C33A cells were transfected with Neo Bam empty, HPV16 E7 and E7 L67R vectors to investigate the mRNA levels of (A) MBD3 and (B) MBD2. (C) Dot Blot experiment was performed to reveal 5hmC levels in C33A cells stably transduced with the Luciferase control and the MBD2 knockdown. Genomic DNA was used at two concentrations (200ng and 50ng). Methylene blue was used as a loading control whereas BSA was used as a negative control. The relative 5hmc level compared to methylene blue stain was plotted on a violin chart. (D) Immunofluorescence validates the increased expression of 5hmC in MBD2 knockdown C33A cells. (E) The global 5hmC levels in CaSki cells transduced with Luciferase and MBD2 knockdown were shown via a dot blot experiment. The bar chart illustrates the relative 5hmC level. (F) Immunofluorescence images verify the elevated expression of 5hmC in MBD2 knockdown CaSki cells. DAPI was used to stain cell nuclei and scale bars indicate 50um. Three independent replicates were used and Plotted values on graphs are the Mean±SEM. The statistical analysis was performed with two-tailed Unpaired T-test with p<0.005 (ns = non-significant, *p<0.05, **p<0.01, ***p<0.001, ****p<0.0001).

### E7 expression transcriptionally mimics MBD2-knockdown C33A cells

Since E7-induced hydroxymethylation is likely mediated through the reduced expression of MBD2 in cells, we hypothesized that the E7-MBD2 interplay could affect transcriptional traits in cancer cells (not only the methylation/ hydroxymethylation status), For this reason, we assumed that E7 expression can mimic the transcriptional profile of genes when MBD2 is knocked down from C33A cells. To check this, we randomly selected genes from different molecular pathways such as the apoptotic, EMT and stemness-related pathways (Fig. 5) and compared their expression when E7 is present in C33A cells versus when MBD2 is knocked down.

**Fig. 5.**
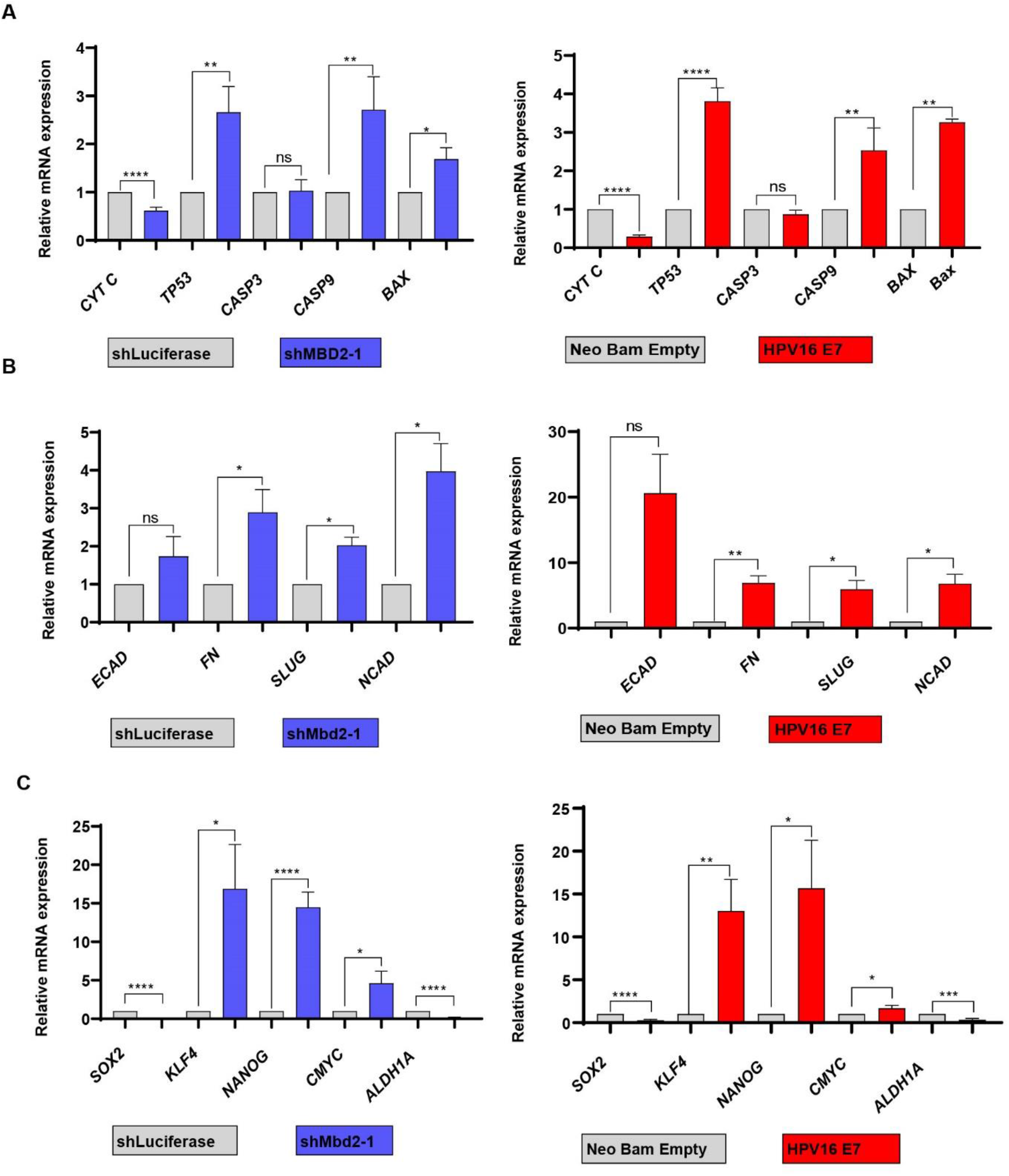
E7 transcriptionally mimics MBD2-knockdown in C33A cells. C33A cells either transduced with the MBD2 knockdown and the Luciferase control or transfected with the Neo Bam empty and HPV16 E7 vectors were collected, and RNA was extracted for qRT-PCR analysis. Genes in the (A) apoptotic, (B) EMT and (C) stemness-associated pathways were investigated. Three independent replicates were used and plotted values on graphs are the Mean±SEM. The statistical analysis was calculated with two-tailed Unpaired T-test with p<0.005 (ns = non-significant, *p<0.05, **p<0.01, ***p<0.001, ****p<0.0001).

Remarkably, we noticed that all 14 genes that we have checked (*CYT C, TP53, CASP3, CASP9, BAX, ECAD, FN, SLUG, NCAD, SOX2, KLF4, NANOG, CMYC, ALDH1A*) display similar trends both in the expression of E7 and in the downregulation of MBD2 in C33A cells. For example, Cytochrome C mRNA expression in MBD2-knockdown cells is reduced compared to the shluciferase control, while a reduced expression was detected in E7-expressing C33A cells (Fig. 5A). Likewise, fibronectin, a key component of the EMT pathway, has an increased mRNA profile when MBD2 is downregulated versus when E7 is expressed in C33A cells (Fig. 5B). Similarly, Sox2 mRNA expression is downregulated in the MBD2-knockdown cells reflecting the same expression profile in E7-expressed C33A cells (Fig. 5C). Nevertheless, we have reported a reduction in the mRNA expression of Oct4 upon MBD2 knockdown in C33A cells (fig. S5A) which this is in contrast of the increased expression of Oct4 in E7-expressing C33A cells [8].

### MBD2 chemical inhibition impairs MBD2 binding on the hOct4 gene and attenuates cell viability

To identify the reason behind the reduced mRNA expression of Oct4 upon MBD2 downregulation, we performed Chromatin immunoprecipitation to determine whether MBD2 binds the hOct4 gene. We observed an enrichment of MBD2 on the hOct4 gene in C33A cells compared to IgG control. However, the MBD2 binding was restricted in the presence of wildtype E7 but not in the presence of the E7 mutant L67R (fig S5A). Hence, we decided to use an MBD2 chemical inhibitor (KCC07), which impairs MBD2 function without affecting MBD2 expression (fig S5), to check whether the MBD2 binding on the hOct4 gene upon the presence of E7 is due to its reduced expression or due to its displacement from Oct4. We used the KCC07 inhibitor at 250nM for 24-, 48- and 72-hours on C33A cells and at 48-hours post-treatment cells were collected for chromatin immunoprecipitation. We noticed that the MBD2 binding on the hOct4 gene was restricted in the presence of the inhibitor both in C33A cells expressing the Neo Bam empty control and the E7 mutant, conditions at which we found previously that MBD2 could bind the hOct4 gene (Fig. 6A). To further characterize the impact of the KCC07 inhibitor on the viability of C33A cells expressing the Neo bam empty control and the wildtype and mutant E7, we treated the cells with 250nM of the inhibitor and checked their cell numbers at 24-, 48- and 72-hours post-treatment. At 48- and 72-hours post-treatment there was a reduction in viability of Empty-expressing and L67R-expressing C33A cells, however this is not the case in E7-expressing C33A cells which remained unresponsive to the inhibitor (Fig. 6B). We also checked the viability of HaCaT keratinocytes and two HPV-positive cervical cancer cells lines (HeLa and CaSki) upon treatment with the MBD2 inhibitor. None of the cell lines had reduced viability when treated with the inhibitor compared to the DMSO control treatment for the 72-hours that we have tested (fig S6A) possibly indicating that normal cells and HPV-positive cancer cells are less sensitive to MBD2 inhibition.

**Fig. 6.**
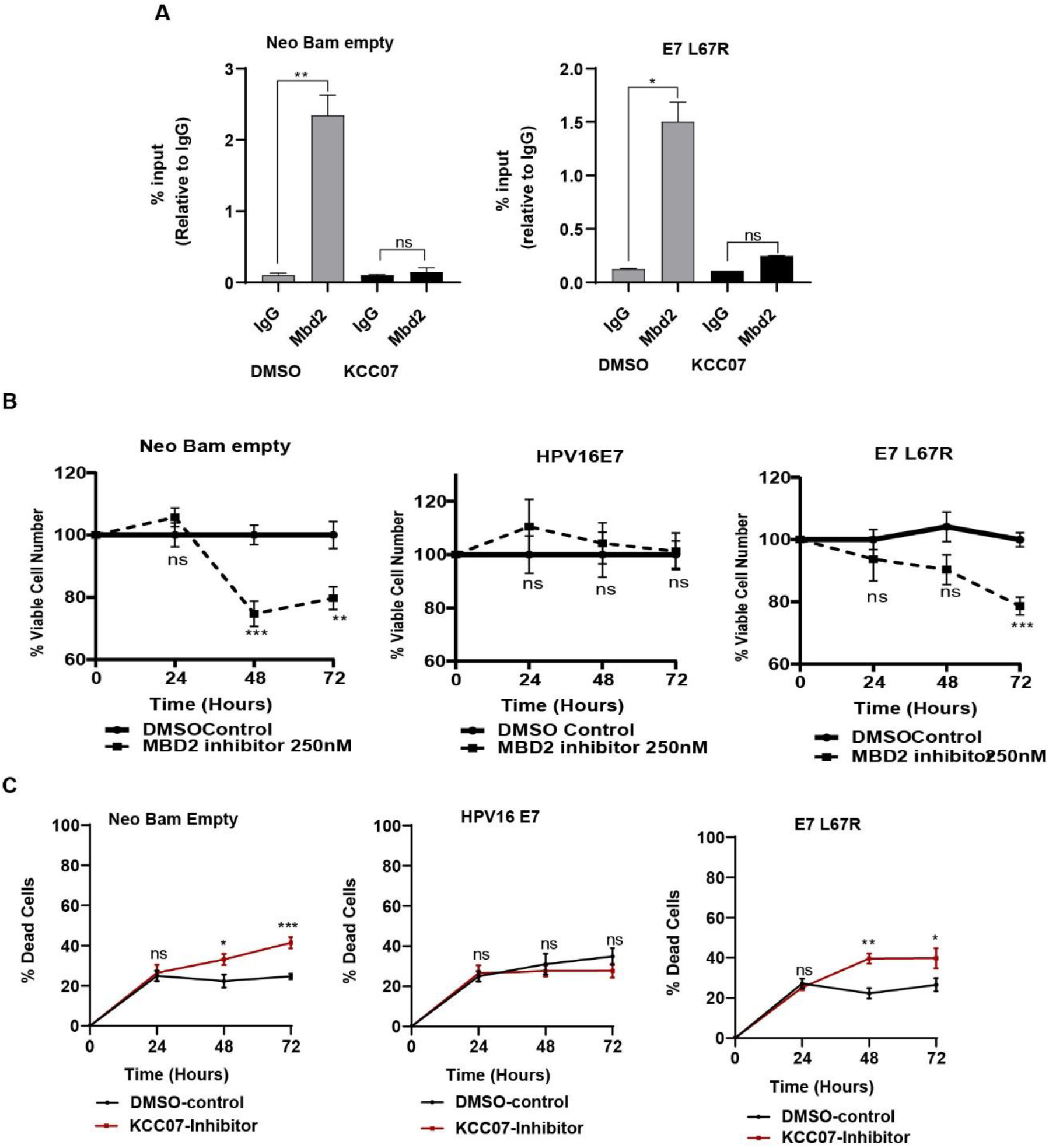
MBD2 inhibition attenuates viability and increases apoptosis in C33A cells not expressing HPV16 E7. (A) The MBD2 inhibitor was used at 250nM concentration to investigate the binding of MBD2 on the hOct4 gene. DMSO was used as a a control for the experiment and 48-hours post application cells were collected for Chromatin immunoprecipitation. (B) KCC07 MBD2 inhibitor was applied on HPV-negative C33A cervical cancer cells transfected with Neo Bam empty, HPV16 E7 and E7 L67R vectors. The inhibitor was applied at 250nM and cell viability was measured for 24-, 48- and 72-hours post-application. DMSO was used as the negative control. (C) Apoptosis was investigated by using the NucGREEN stain and apoptotic (green) cells were visualized under the microscope. Three independent replicates were used and plotted values on graphs are the Mean±SEM. The statistical analysis was calculated with two-tailed Unpaired T-test with p<0.005 (ns = non-significant, *p<0.05, **p<0.01, ***p<0.001, ****p<0.0001).

To examine the mechanism behind the reduced viability in C33A cells upon treatment with KCC07, we asked whether these cells were subjected to cell death. For this reason, we used the NucGreen stain which gets incorporated in the nucleus of apoptotic cells due to the fractured cellular membranes. We quantified the number of green nuclei in KCC07-treated versus DMSO-treated cells and we detected an increase in the apoptotic cells at 48- and 72-hours post-treatment in Empty- and L67R-expressing C33A cells (not in E7-expressing cells) (Fig. 6C). Additionally, we checked the mRNA profile of apoptotic genes in Empty-, E7- and L67R-expressing C33A for 24-, 48- and 72-hours post-treatment. We observed that the expression of these genes in E7-expressing cells did not change upon treatment with KCC07, while Empty- and L67R-expressing cells have a deregulated expression of *BCL2*, *BAX*, *CASP3* and *CYT-C* further validating the increase in apoptosis we observed upon KCC07 treatment (fig S6B-D). An RT-PCR analysis was performed to verify the expression of wildtype and mutant E7 in C33A cells (fig S6E).

## Discussion

The upregulation of Oct4 is reported in many cancer types including cervical cancer [2, 4–7]. We have previously shown that HPV-driven cervical cancers express higher Oct4 levels compared to non-HPV-driven cervical tumors. The viral oncogene E7 accounts for the upregulation of Oct4 in HPV-driven cervical tumors as it interacts with Oct4 at its CR3 domain [8]. To further characterize the mechanism by which Oct4 is re-expressed in cervical cancer, we profiled the Oct4 interactome in C33A cervical cancer cells. Even though there is increasing evidence implicating Oct4 in several cancer types, we have identified no other reports of the Oct4 interaction network in cancer presumable due to the relatively low levels of Oct4. To circumvent expression issues, we overexpressed Oct4 in cells and performed Affinity Purification-Mass Spectrometry to identify the network of proteins interacting with Oct4 in cervical cancer cells. Parallel proteomic and bioinformatics analyses revealed the NuRD complex as one of the complexes that interact and regulate Oct4. Nevertheless, the exact mechanism by which this complex or its individual components regulate Oct4 expression during cervical tumorigenesis remains unclear. In this manuscript, we revealed certain components of the NuRD complex to be associated with Oct4, while E7 expression affects the interaction between Oct4 and NuRD components. Notably we report here that Oct4 interacts with two variants of the NuRD complex: in the absence of E7, Oct4 mainly interacts with MBD2-NuRD whereas in the presence of E7, Oct4 interacts with MBD3-NuRD. These two MBD proteins are mutually exclusive in the NuRD complex [49] further validating our data. However, we do not know whether the differential binding of Oct4 with MBD2 and MBD3 in the presence/absence of E7 is due to stochiometric issues or it stems from a mechanism that has not yet been identified.

Evidence indicates a stronger binding of MBD2 to hypermethylated promoters, as opposed to MBD3 which has a less tight attachment [50, 51]. MBD3 binds strongly to unmethylated or hydroxymethylated regions on the DNA such as enhancers and gene bodies [53]. DNA hydroxymethylation is considered as the first derivative of the active demethylation process and is considerably associated with transcriptional activation contrary to the transcriptional repression mediated by DNA methylation. Hydroxymethylation is mediated by the TET enzymes that catalyse the oxidation of 5mC to 5hmC. The TET enzymes further convert 5hmc to 5fC and 5caC where the conversion of 5caC to unmethylated cytosines is facilitated by the TDG enzyme (fig S2A). The process by which TDG removes oxidized bases to maintain the integrity of the DNA is called Base Excision repair (BER). It is reported that the BER mechanism, mediated via the interaction between TET1 and TDG, is linked to the active demethylation cycle [54] further reinforcing our data. Upon the presence of E7 there is an increase in the mRNA and protein expression of TET1 and TDG, promoting the increase in global hydroxymethylation (Fig. 2, A and B) (fig S2B).

We propose a fundamental role of E7 in the re-expression of Oct4 in cervical cancer cells (fig S7) and in the onset and progression of the demethylation cycle in cervical cancer and via the re-expression or activation of the TET1 enzyme (Fig. 2, A and B). In somatic cell reprogramming, TET1 can substitute Oct4 for the formation of induced pluripotent stem cells via demethylating and re-expressing Oct4 [58]. TET1 induces 5hmC enrichment throughout the process of reprogramming showing that the methylation/demethylation status is important for this process demonstrating that TET1 mediated regulation of Oct4 expression may be relevant in contexts other than carcinogenesis.

E7 increases TET1 expression and elevates total 5hmC, nevertheless, our immunofluorescence data do not show major differences in total methylation levels (Fig. 2D). Since these are not accurate measures of local transcriptional regulation, we determined the methylation-demethylation balance locally, by checking 5mC and 5hmC levels on the hOct4 gene in the presence or absence of E7. Randomly selected loci on the hOct4 gene were designed (areas upstream the promoter, areas spanning the promoter, areas around TSS and areas downstream TSS) (fig S3C) to test their methylation or hydroxymethylation status. In the absence of E7, all loci tested on the hOct4 gene were found to be methylated probably suggesting that the Oct4 gene in C33A cells is methylated. However, upon the presence of E7 previously methylated sites on the hOct4 gene were hydroxymethylated (Fig. 3B) possibly proposing an underlying mechanism of Oct4 re-expression during cervical carcinogenesis. Notably, not all loci tested were converted to a state of hydroxymethylation, but these are areas upstream or downstream the promoter which support the fact that hydroxymethylation happens mostly on gene bodies and enhancers, not promoters [59].

Apart from the re-expression of Oct4 in virally induced hepatocellular carcinoma via the activation of IL-6 [60], this is the first time we introduce a potential mechanism of Oct4 re-activation during the carcinogenic process.

In this manuscript we demonstrate that the E7-mediated increase in 5hmC is more likely linked to a reduction in MBD2 levels rather than MBD3. E7-expressing C33A cells do not show a change in the MBD3 mRNA expression, although we reported a decrease in the MBD2 levels (Fig. 4B). According to Ludwig et al, MBD2 guards the DNA from TET1-induced oxidation [61] supporting our findings that upon E7-mediated downregulation of MBD2 there is an increase in hydroxymethylation levels (Fig. 4). We further characterized the molecular interplay between E7 and MBD2 in cervical cancer cells, by checking the mRNA expression profile of 15 randomly selected genes upon E7 expression and MBD2 knockdown in C33A cells. 14 out of 15 genes were expressed in the same way in E7-expressing and MBD2-knockdown C33A cells (Fig. 5) signifying a transcriptional mimicry between E7 and MBD2 downregulation. Nonetheless, Oct4 downregulation upon MBD2 knockdown (fig S5A) does not match the Oct4 upregulation we see in E7-expressing cells [8]. To understand this disparity in Oct4 expression we examined whether Oct4 is regulated by both E7 and MBD2 via chromatin immunoprecipitation. Remarkably we noticed that E7 restricts the binding of MBD2 on Oct4 while when E7 is absent or mutated in C33A cells, MBD2 can bind to Oct4 promoter regions (fig S5B). However, we could not effectively identify whether the restricted binding of MBD2 on Oct4 in the presence of E7 is due to the downregulated MBD2 expression in E7-expressing cells or a due to a competitive disadvantage between E7 and MBD2 to bind Oct4. Since L67R-expressing C33A cells yield a reduced mRNA expression of MBD2 (Fig. 4B) but L67R-expressing C33A do not impair the binding of MBD2 on Oct4 (fig S5B), we hypothesized that it is not the decreased levels of MBD2 that impact the restricted binding on the Oct4 gene but there must be a mechanism that it is yet unknown leading to a competitive binding between MBD2 and E7 on the Oct4 gene.

To further support our hypothesis, we used an MBD2 inhibitor (KCC07), which was previously used to treat Medulloblastoma [62]. This inhibitor impairs the binding of MBD2 on the DNA without affecting its expression. We provided substantial support that upon treatment with KCC07 in C33A cells, MBD2 could not bind Oct4 compared to the DMSO control treatment where the MBD2 binding was evidenced. The MBD2 inhibitor reduced the viability of C33A cells and increased apoptosis (similar to effects seen in medulloblastoma cells). E7-expressing cells (C33A cells ectopically expressing wild type E7, or HPV-positive cervical cancer cells) were less sensitive to MBD2 inhibition overall suggesting that switching to alternate MBD variants is a broader mechanism of transcriptional regulation during HPV-driven carcinogenesis.

We report here a new mechanism by which E7 regulates transcription, providing a broader paradigm by which Oct4 and other silenced genes are re-expressed during carcinogenesis. The importance of this mechanism during cervical carcinogenesis may be more critical during early stages of carcinogenesis, as others have reported elevated levels of TET1 during cervical premalignancy [63, 64]. Even thought a lot of studies suggest a substantial increase in the global methylation of HPV-positive tumors [65–69], bisulfite sequencing is not a robust method in differentiating between methylation and hydroxymethylation. Moreover, aberrant DNA methylation/demethylation may vary among specific loci, despite overall patterns. In light of our findings, we propose that a reassessment of DNA methylation status in HPV-related cancers using modalities which more effectively differentiate between methylation/hydroxymethylation, may be warranted.

## Materials and Methods

### Cell Lines and Culture

CaSki and C33A cervical cancer cells were purchased from ATCC and maintained in DMEM and MEM (mixed with 1% L-glutamine) respectively. 293T epithelia cells (ATCC) and HaCaT immortalized keratinocytes (CLS) were maintained in DMEM. All culture media were supplemented with 1% Penicillin/streptomycin (P/S) and 10% Fetal Bovine Serum (FBS). For the transfection experiments, C33A and HaCaT cells were placed at a density of 3x10^5^ and 5x10^5^ respectively. 24-hours post-placing cells were transfected with Fugene transfection reagent. For the transduction experiments 293T cells were placed at a 1x10^6^ density. Mammalian and retroviral plasmids were used as shown in Table S3. 48-, 72- and 96-hours post-transfection, the retrovirus was collected and applied in C33A, CaSki and HaCaT cells mixed with 1ug/ml Polybrene. Stable cells were selected with specific antibiotics as shown in Table S3.

### RNA extraction, cDNA synthesis and PCR

Cells used for RNA extraction were isolated with the Trizol method. To remove DNA impurities from RNA extracts, the AMBION-DNA free kit was used and 300ng of cDNA was synthesized with the iSCRIPT cDNA synthesis kit. For RT-PCR (Reverse-Transcription-Polymerase chain Reaction) the KapaTaq PCR kit was used whereas qPCR (Real time-PCR) was achieved by KAPA SYBR FAST qPCR Master Mix (2X) kit according to the manufacturers’ guidelines. The primer sequences used are cited in Table S1. For each gene, the average C(t) value was determined and was normalised to housekeeping (*GAPDH* or *ACTIN*) genes. Unpaired t-test (two-tailed) was used to calculate statistical significance.

### 3’-mRNA Quant Sequencing

C33A cells were transfected with i) Oct4+Neo Bam empty and ii) Oct4+HPV16 E7 vectors. 48-hours post-transfection the cells were collected for RNA extraction with The RNeasy mini kit by Qiagen and the expression of Oct4 and E7 was validated with RT-PCR. For the Quant-sequencing and analysis of the samples, we sent the RNA to B.S.R.C. Fleming Institute in Greece. The differentially expressed gene (DEG) analysis was performed with the PANDORA algorithm. Following normalisation with the metaseqr2 pipeline and filtering (to remove artefacts) we selected and identified genes with a p<0.05 value and a fold change greater than 2 (FC>2) [70]. For the validation of the DEGs we used primers targeting some highly upregulated (*THSB1, C-MYC, RGS4, ANKRD1, BZW2, PTPRC*) and highly downregulated (*GLS2, CHDR, ZNF483*) genes.

### Computational analysis to identify Oct4 and E7 PPIs

To identify Oct4 and E7 PPI we searched the Universal Protein Resource (UniProt) [71] using the queries `<gene:pou5f1 OR gene:oct4 OR gene:oct3 OR name:“pou domain class 5 transcription factor 1>` and `<e7 organism:human AND gene:e7>`. The query keywords for Oct4 retrieved 13 protein entries from 9 different species, while E7 query keywords retrieved 66 protein entries from 66 different HPV strains. The identifiers gathered were used as query keywords for the collection of PPI data of Oct4 and E7 proteins. Relevant data were downloaded from IntAct (version 4.2.16) [72], BioGRID (release 3.5.186, 25/05/2020) [43] and Agile Protein Interactomes Dataserver (APID) (version January 2019) [42]. The datasets downloaded from APID served as templates where additional PPI data were added by searching manually literature papers. All available PPI datasets as gathered from APID were manually examined thoroughly in order to minimize the possibility of having missing or inaccurate values. By using UniProt’s Retrieve/ID mapping tool every protein interactor was mapped to its current UniProt identifier as well as its HGNC (HUGO Gene Nomenclature Committee) approved gene name. Regarding the experimental methods used for the identification of each interaction we used the Ontology Lookup Service (OLS) [44]. Specifically, Molecular Interactions Controlled Vocabulary (MICV) Ontology terms were gathered from OLS to describe in more detail the entries regarding the experimental/computational methods that were used to identify the PPIs. Homology of Oct4 proteins and their interactors was evaluated by exploiting Phylogenomic databases such as KEGG Orthology (KO) [73], OrthoDB [74] and InParanoid [75], which provide information about the existence of orthologous proteins between different species. Taking into consideration the importance of E7 oncoprotein and its role in the viral life cycle all available E7 proteins of HPV strains were accounted as homologs. For each PPI a score was calculated based on a simplified form of IntAct’s scoring methodology (MIscore) as described in **Equation 1** [76]. For each PPI there is information about the method used for its identification, the type of that method and the publication from where the interaction was extracted. For each method and method type ontology term, a specific score is assigned.

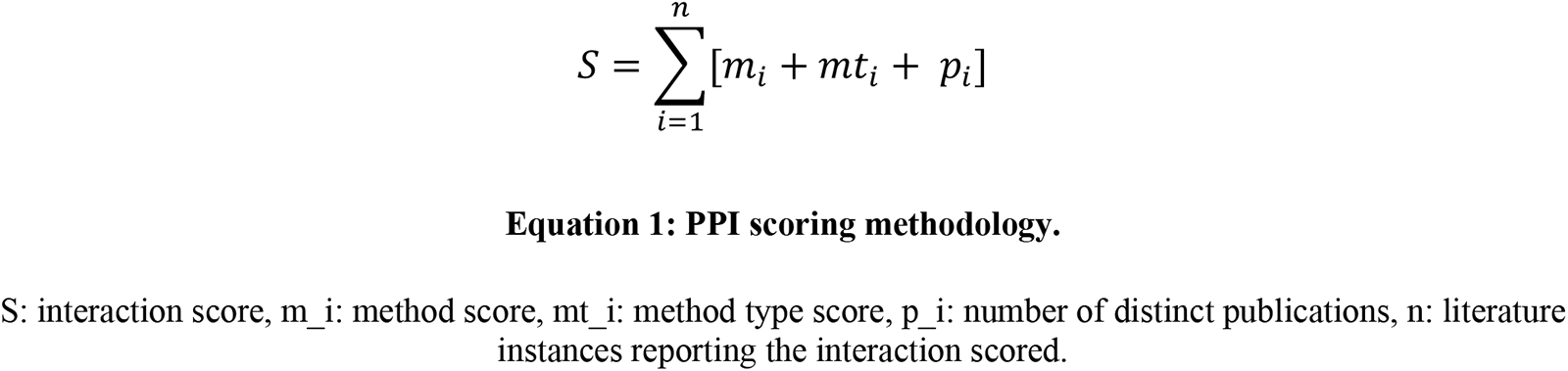

Method and method-type scores are summed up for each interaction and if there is more than one indication for that interaction the score is the sum of the individual observations. Then, to that score the number of distinct publications referring to the interaction is added, and the score is normalized based on all entries in the database. This formulation was used to score all interaction pairs for hOct4 and HPV16 E7.

In order to integrate Oct4/E7 interactors from other species/strains we use a slightly altered scoring methodology, which also takes into consideration the evolutionary distance as encoded in the sequence similarity to their human/HPV16 counterparts. In more details, when summing the method and method-type scores (as in Equation 1) we weight their contribution by the percentage identity to the source interactor (hOct4 or HPV16 E7) as determined by BlastP (run with default settings against the “Non-redundant protein sequences (nr)” database at the NCBI; accessed November 2020) (Equation 2). This way interologs from more distantly related species are downweighed, which is a conservative approach to integrating this information to the human PPI network without introducing many false positives.

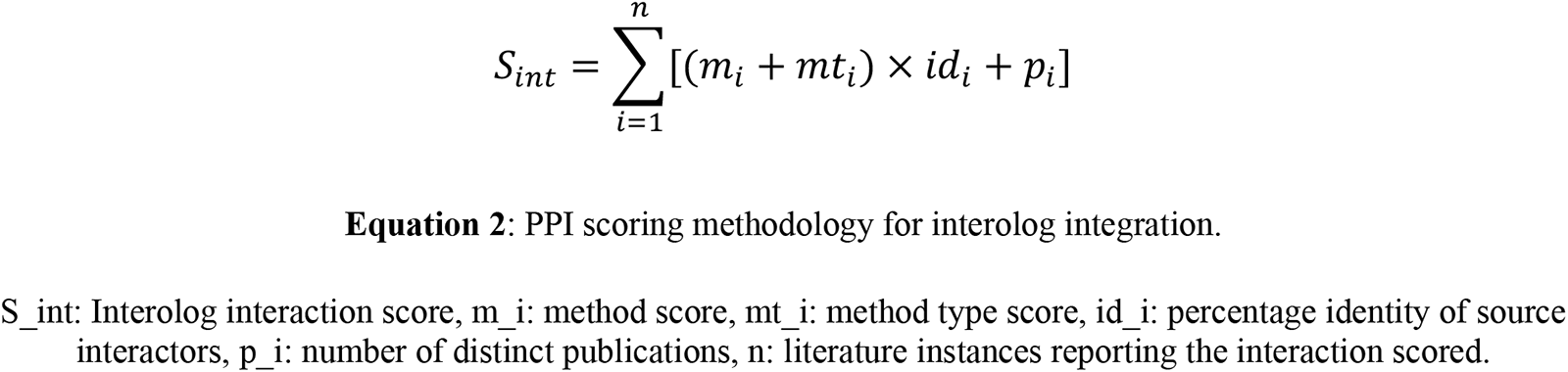

The scores of the non-redundant PPIs (including interologs) are then normalized for computing a final score using the Min-Max normalization method: 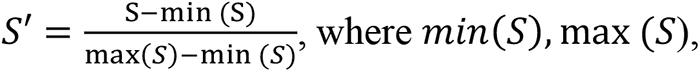 correspond to the minimum and maximum values of observed scores in the Oct4 and E7 dataset respectively. Notable the normalized score lies in the [0,1] range. Normalized PPI scores are further binned into 3 categorical ranges as follows: weak, when *S*′ ∈ [0,0.333); medium, when *S*′ ∈ [0.333, 0.666); strong, when *S*′ ∈ [0.666,1]. Finally, the PPIs are used to construct the interactor networks of Oct4 and E7. The data used and the Python code developed for our analysis can be found in https://github.com/MariosEft97/bachelor_thesis.

### Protein extraction and Western Blotting

Cells collected for Western blotting were lysed with RIPA cell lysis buffer (150mM NaCl, 5mM EDTA, 50mM Tris-HCL,1% TritonX-100, 0.1% SDS and 0.5% Sodium Deoxycholate) supplemented with Protease/Phosphatase inhibitors. The concentration of protein samples was quantified and normalised by Bradford. 80ug of protein extracts were used and loaded onto a SDS-PAGE gel for electrophoresis. Proteins were then transferred onto a nitrocellulose membrane by using the Wet transfer apparatus. The membranes were blocked with 5% BSA in TBS-Tween blocking buffer for 1 hour at room temperature. The membranes were then incubated overnight at 4°C with primary antibodies as mentioned in Table S4 and secondary antibodies conjugated with HRP were added and incubated at room temperature for 1 hour. Protein expression was then viewed with ECL reagents by using G-box imager.

### Co-Immunoprecipitation

C33A cells were transfected with Oct4, Oct4+HPV16E7 and Oct4+E7 L67R vectors and 48-hours post transfection, cells were collected for immunoprecipitation. Protein extraction was achieved as mentioned above using ice-cold RIPA lysis buffer supplemented with protease inhibitors. Protein samples were pre-cleared with 1:1 Sepharose beads in RIPA buffer for 1 hour under low agitation. Following centrifugation at 120000g for 30 seconds to remove the beads, the protein lysates were separated for the a) input control sample, b) IgG negative control and c) for immunoprecipitating Oct4. The primary antibodies were incubated with the protein lysates overnight at 4°C under low agitation. Next day the slurry (Sepharose beads in RIPA buffer) was added to the antibody-lysate mixture and incubated for 3 hours at 4°C under low agitation. The slurry and antibodies were removed following boiling the samples at 95°C for 10 minutes and then the samples were loaded onto SDS-PAGE gels for electrophoresis. Then Oct4-protein interactions were visualized via western blotting.

### Mass Spectrometry

C33A cells were used for the Mass Spectrometry experiment to identify the Oct4 interactome. To avoid specificity issues and variability in detection limits with the Mass spectrometry due to the low abundance of Oct4 in C33A, we transfected the cells with an Oct4 vector (Table S3). Following successful transfection, immunoprecipitation was performed using an Oct4 antibody following a pre-clearing step with 1:1 Sepharose beads diluted in RIPA buffer. The negative control used was an IgG antibody to ensure that the immunoprecipitation of Oct4 is specific to the Oct4 antibody. Additional control for the experiment was a stable Oct4-knockdown C33A cell-line which we already established [8]. The Mass spectrometry was performed, and the raw data and an initial analysis were provided by the EMBL Proteomics Facility. Interactors were identified with the R-software and the duplicated interactors were removed. Data were normalised by using the vsn package [77]. The peptide reads were classified as **hits (when** *false discovery rate [FDR]< 0.05 and a fold change of at least 2-fold*) and **candidates** (*FDR < 0.2 and a fold change of at least 1.5-fold*) with LIMMA analysis. The top interactors were validated by Western Blot.

### Enrichment Analysis

Enrichment analyses to identify gene pathways and protein complexes from the Quant-seq and Mass Spectrometry experiments were achieved by using the online software Enrichr [78–80].

### DNA extraction and Dot Blotting

C33A, CaSki and HaCaT cells were collected for the Dot blot experiments at a density of 6x10^6^ cells. Cell pellets were treated as indicated by the DNAeasy Blood and Tissue kit by Qiagen. Eluted genomic DNA was quantified by nanodrop and boiled to 95 °C for 10 minutes to allow denaturation and then a volume of 1.5ul of the samples was added on a nitrocellulose membrane in the form of dots. Membranes were then blocked in 5% BSA mixed in TBS-Tween for 1 hour at room temperature and then were incubated with primary antibodies Table S3 for 1 hour at room temperature. HRP-conjugated antibodies were added to the membranes for 1 hour at room temperature and then dots were visualized with ECL reagents at a G-box imager.

### Immunofluorescence

Cervical cancer cells (C33A & CaSki) and immortalized keratinocytes (HaCaT) were seeded on coverslips at a density of 1x10^5^ and 5x10^5^ respectively. 24-hours post-placing the cells were washed with sterile 1X-PBS and fixed with 4% paraformaldehyde. Permeabilisation was achieved with 0.2% Triton X-100 and 5% BSA blocking buffer was used on the coverslips for 30 minutes. Primary antibodies Table S3 were diluted in blocking buffer and added to the cells overnight at 4°C. FITC anti-rabbit and Alexa-594 anti-mouse secondary antibodies were added and incubated at room temperature for an hour. Cells were then viewed with a Fluorescent Zeiss microscope. Vectashield antifade mounting medium with DAPI (Vector H-1200) was used for mounting the slides and staining cells nuclei. Images were processed and analysed by using the ImageJ software [81].

### NucGreen Dead Stain

For imaging and quantifying dead cells, the NucGreen Dead-488 Ready Probes Reagent from Invitrogen was used. C33A cells transfected with Oct4, HPV16 E7 and E7 L67R vectors were treated with 250nM KCC07 MBD2 inhibitor or the DMSO control. At 24-, 48- and 72-hours post treatment the cells were treated with the NucGreen Reagents as per the manufacturer protocol and then visualized with a Fluorescent Zeiss Microscope and analysed with ImageJ.

### ChIP-qPCR

Cervical cancer C33A cells were transfected with i) Neo-Bam empty, ii) HPV16E7 and iii) E7 L67R vectors. 1x10^7^ cells were used and DNA-protein complexes were cross-linked with 1% Formaldehyde for 10 minutes and 125mM Glycine was added for 5 minutes for the quenching process. Cells were washed with ice-cold sterile 1xPBS and collected with centrifugation at 2000rpm for 5 mins at 4 °C. ChIP lysis buffer (50mM HEPES-KOH pH7.5, 140mM NaCl, 1mM EDTA pH8, 1% Triton X-100, 0.1% Sodium Deoxycholate, 0.1% SDS, Protease Inhibitors added last) was added to the cell pellet for 10 minutes (on ice) and sonication to allow chromatin shearing was performed with the Diagenode Bioruptor^TM^ UCD-200 Sonicator to achieve a chromatin length of around 100-300bp (Sonication pulses 7seconds on nd 7 seconds off for a period of ten minutes). 15ug of the sheared chromatin was used for the immunoprecipitation step with primary antibodies (table S3) incubated for 1 hour at 4 °C and then 50ul sepharose beads primed with 75ng/ul herring Sperm and 0,1ug/ul BSA were added. The immunoprecipitated DNA was treated with 5M NaCL, 10mg/ml RNase A and incubated overnight at 65 °C. Proteinase K 20mg/ml was added and incubated at 65 for 1 hour and DNA was purified with PCR purification Kit from Qiagen. Then, the purified DNA was used to perform qPCR by using primers (table S1) for specific loci on the hOct4 gene.

### meDIP-ChIP and hmeDIP-ChIP

Cervical cancer C33A cells were transfected with i) cmv-Neo-Bam empty and ii) cmv-HPV16E7 vectors. 48-hours post transfection cells were collected and fixed as mentioned above. Sonication was achieved by using the Diagenode Bioruptor^TM^ UCD-200 Sonicator to reach a chromatin length between 100-300bp. 25ug of sheared chromatin was used for meDIP and hmeDIP kits from Abcam to examine methylation and hydroxymethylation on the hOct4 gene respectively. The guidelines from the manufacturer company were followed and the eluted DNA was quantified using a nanodrop. Primers targeting different regions on the hOct4 gene were used for qPCR (table S2).

### Statistical Analyses

Statistical analyses were plotted and quantified using the GraphPad Prism v.6.0 (La Jolla, CA). All the experiments were performed using at least three biological replicates. Statistical significance was calculated at p<0.05.

Parts of the figures were generated using the Biorender.com.

## Acknowledgments

We thank members of the KS lab for their useful feedback for this manuscript.

## Author Contributions

Conceptualization: KS, VP, TP

Methodology: TP, ME, EP

Investigation: TP, ME, EP

Analysis: TP, ME, EP, KS, VP

Supervision: KS, VP

Writing—original draft: TP, KS

Writing—review & editing: TP, MS, KS, VP

## Data and materials availability

Sequencing: The data for this study have been deposited in the database UCSC using the following URL: http://epigenomics.fleming.gr/~alexandros/strati_quantseq/hub.txt

Code for computational analysis: The data for this study have been deposited in the database GitHub using the following URL: https://github.com/MariosEft97/bachelor_thesis

Mass Spectrometry: All of the raw Mass Spectrometry data are found on the ProteomeXchange Consortium via MassIVE with the description MSV000091651.

