## Supplemental files for "A paradigm for post-embryonic Oct4 re-expression: E7-induced hydroxymethylation regulates Oct4 expression in cervical cancer"

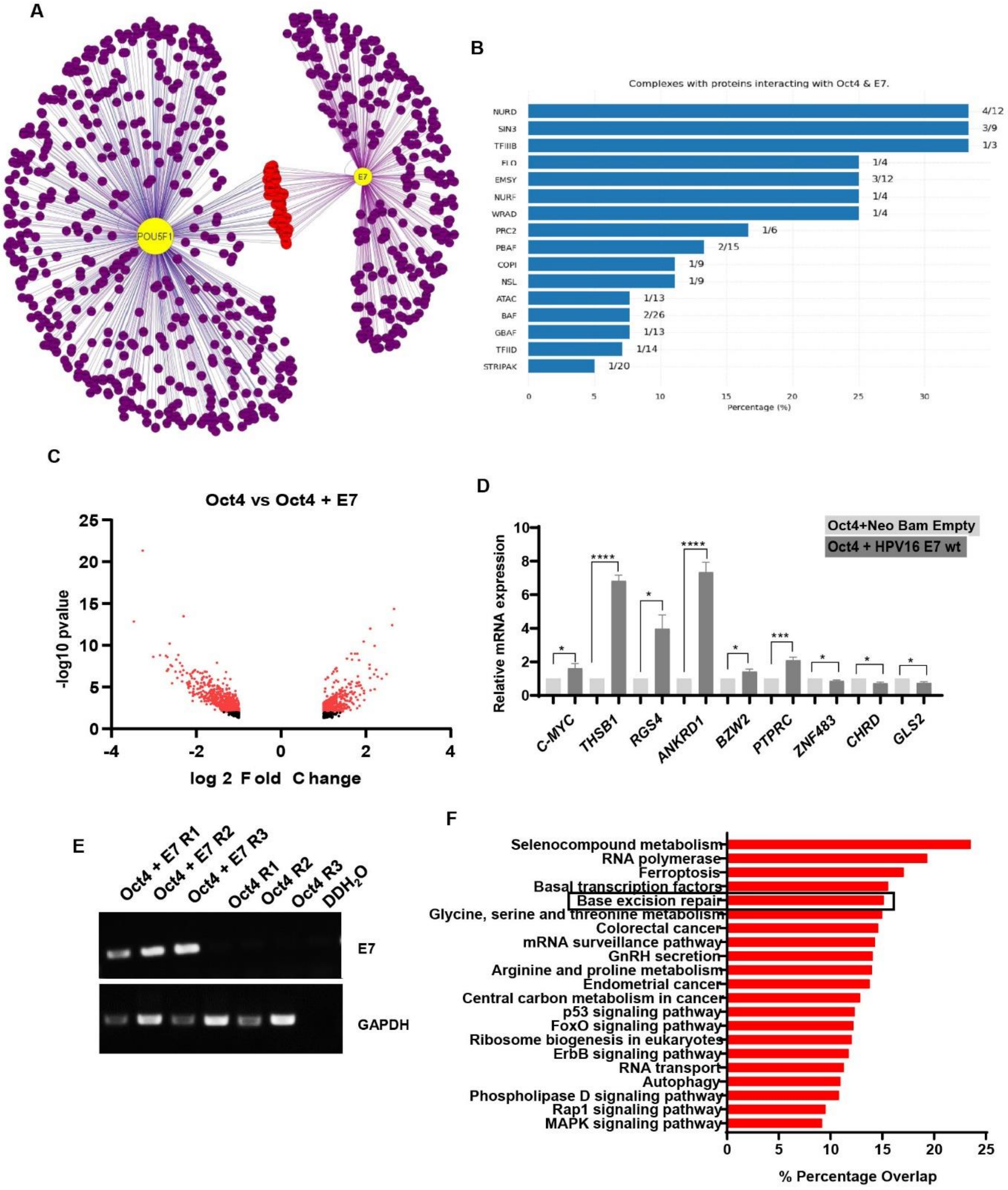


**Fig. S1.**

**Proteins and genes regulated by Oct4 and E7 in cervical cancer cells.** (A) Computational analysis demonstrating the Oct4 & E7 protein interaction partners. Oct4 and E7 are illustrated in yellow while their interactors are colored purple. Common interactors of the two proteins are shown in red (Common interactors are: UBR4, UBR5, KCMF1, RBM4, DNAJA3, GATAD2B, C14orf93, COPA, SAP30BP, AP2M1, CSNK2A1, WDR5, CCT3, PAK4, MYC, PKM, ACOT9, PPP2R1A, CREBBP, FUBP3, ZGPAT, CHD4, ELOC, HDAC1, HDAC2, TBP, CDK1, KDM1A, NUMA1, PML, SMAD3, SMARCA4, TUBG1, RBBP7, SRP9, PSMB2, 27 SNRPB2, TCF20, SMARCC2, PPP1CA, TAOK1). There are 834 nodes and 877 edges and edge color corresponds to the score-based binning each the interaction (Weak: purple, Medium: blue, Strong: black). (B) Protein complexes interacting with Oct4 and E7 are revealed. The higher the percentage of individual proteins participating in a complex and interact with Oct4 and E7 are shown on the top part of the graph. (C) Volcano plot illustrating differentially expressed genes (DEGs) in Oct4-expressing and Oct4+E7-expressing C33A cells as shown by the Quant-Seq analysis. (D) Validation of the Quant-Seq with qRT-PCR by using some top regulated genes that are either upregulated or downregulated in Oct4 versus Oct4+E7 condition. (E) RT-PCR was conducted to show the successful transfection of E7 in C33A cells. Gapdh was used as the loading control. (F) Bar chart showing the pathways where DEGs belong to according to the Enrichr software. Three independent replicates were used for the Quant-seq experiment, and the statistical analysis was performed with two-tailed Unpaired T-test with $p<0.05$ (ns = non-significant, *$p<0.05$, **$p<0.01$, ***$p<0.001$, ****$p<0.0001$) and plotted values in the graph are mean±SEM.

A

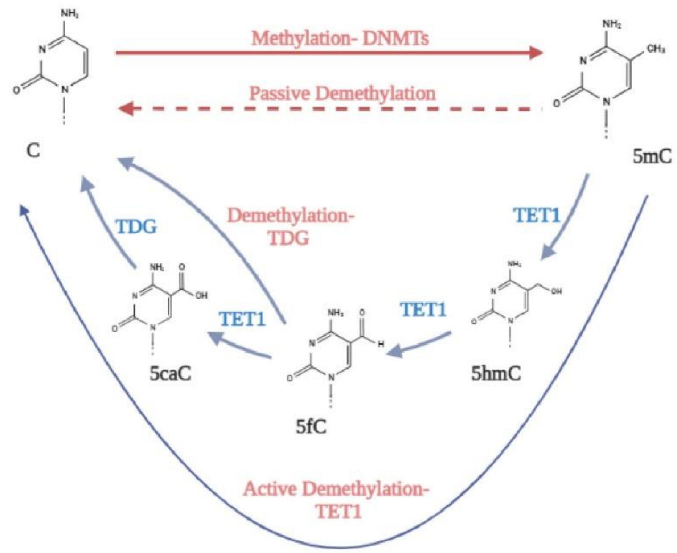


B

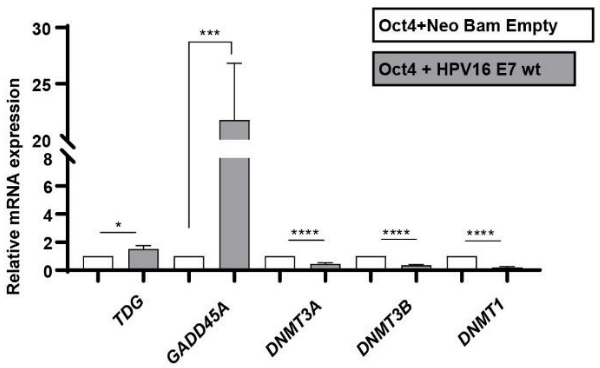


**Fig. S2.**

**E7 regulates enzymes involved in the active demethylation cycle.** Diagrammatic representation of the methylation-demethylation cycle. DNMT enzymes add methyl groups on cytosine residues on their 5-carbon position of the nucleosomes (5mC). The TET1 enzyme catalyses the oxidation of 5-methylcytosine to 5-hydroxymethylcytosine (5hmC) and to subsequent conversions to 5fC and 5caC. TDG is an enzyme in the base excision repair (BER) mechanism converting 5caC back to the unmethylated cytosine. (B) mRNA expression of genes involved in the demethylation process as indicated by the Quant-Seq data. Three independent replicates were used for the Quant-seq experiment, and the statistical analysis was performed with two-tailed Unpaired T-test with $p<0.05$ (ns = non-significant, *$p<0.05$, **$p<0.01$, ***$p<0.001$, ****$p<0.0001$) and mean±SEM.

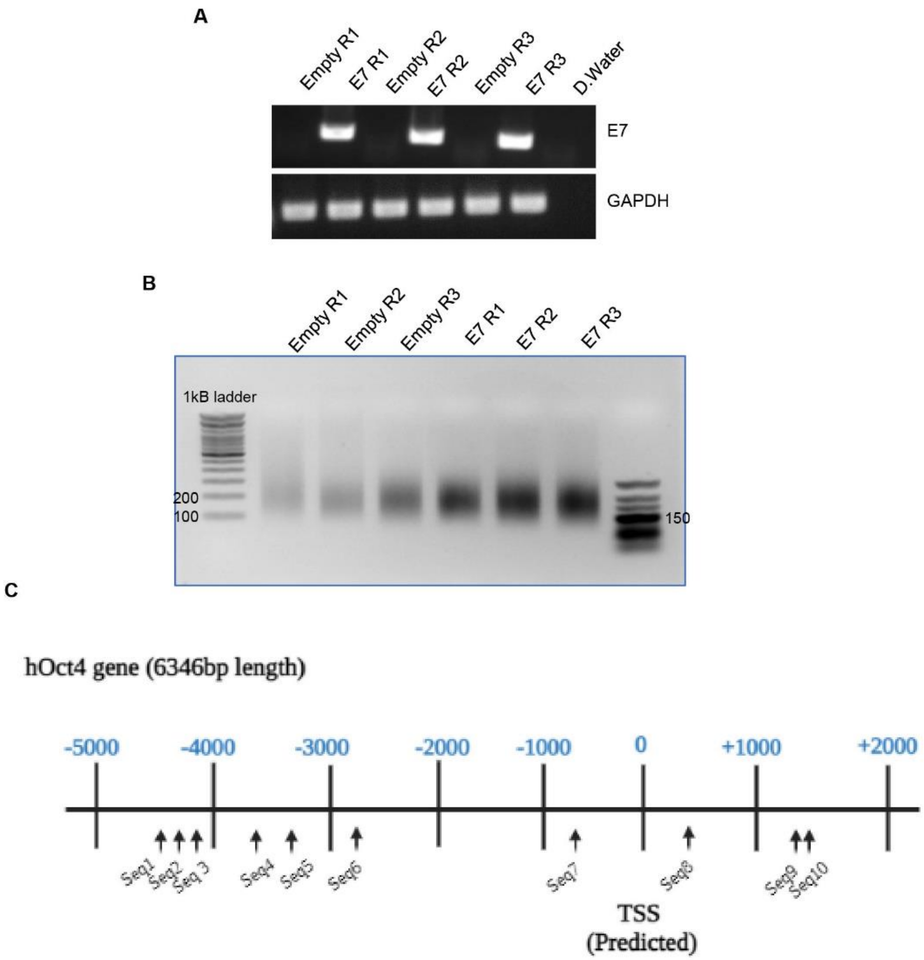


**Fig. S3.**

**Validation of Chromatin immunoprecipitation experiments and sequences binding the hOct4 gene.** C33A cells were used for chromatin immunoprecipitation experiments (Methylated DNA Immunoprecipitation and Hydroxymethylated DNA immunoprecipitation) and were transected with Neo Bam empty and HPV16E7 vectors. (A) RT-PCR was performed to verify the transfection of E7 in C33A cells and GAPDH was used as a control. (B) Sheared chromatin was visualized in an agarose gel to check its size. Three independent replicates were used. (C) Diagrammatic representation displaying the location of sequences used for Chromatin Immunoprecipitation targeting the human Oct4 gene (complementary strand).

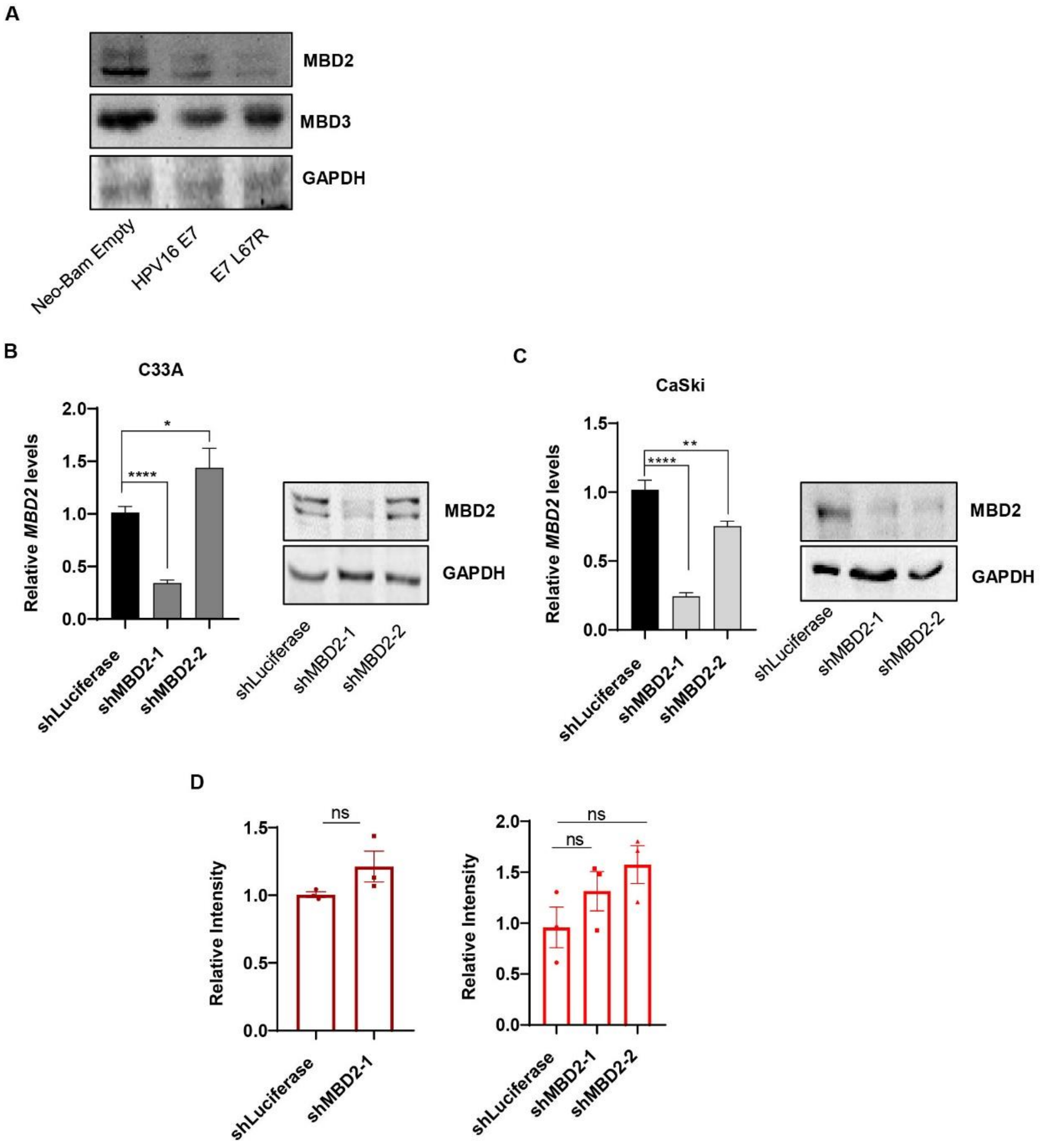


**Fig. S4.**
**MBD2 knockdown in cervical cancer cells.** (A) C33A cells transfected with the Neo Bam Empty, HPV16 E7 and E7 L67R vectors were assessed for MBD2 and MBD3 protein levels. Validation for the MBD2 knockdown in (B) C33A and (C) CaSki cells at the mRNA and protein level. GAPDH was used as a control. (D) Quantification of the relative intensity of 5hmC levels in C33A and CaSki cells stably expressing the shLuciferase control and the MBD2 knockdown was plotted. Three independent replicates were used and Plotted values on graphs are the Mean±SEM. The statistical analysis was performed with two-tailed Unpaired T-test with $p<0.05$ (ns = non-significant, *$p<0.05$, **$p<0.01$, ***$p<0.001$, ****$p<0.0001$).

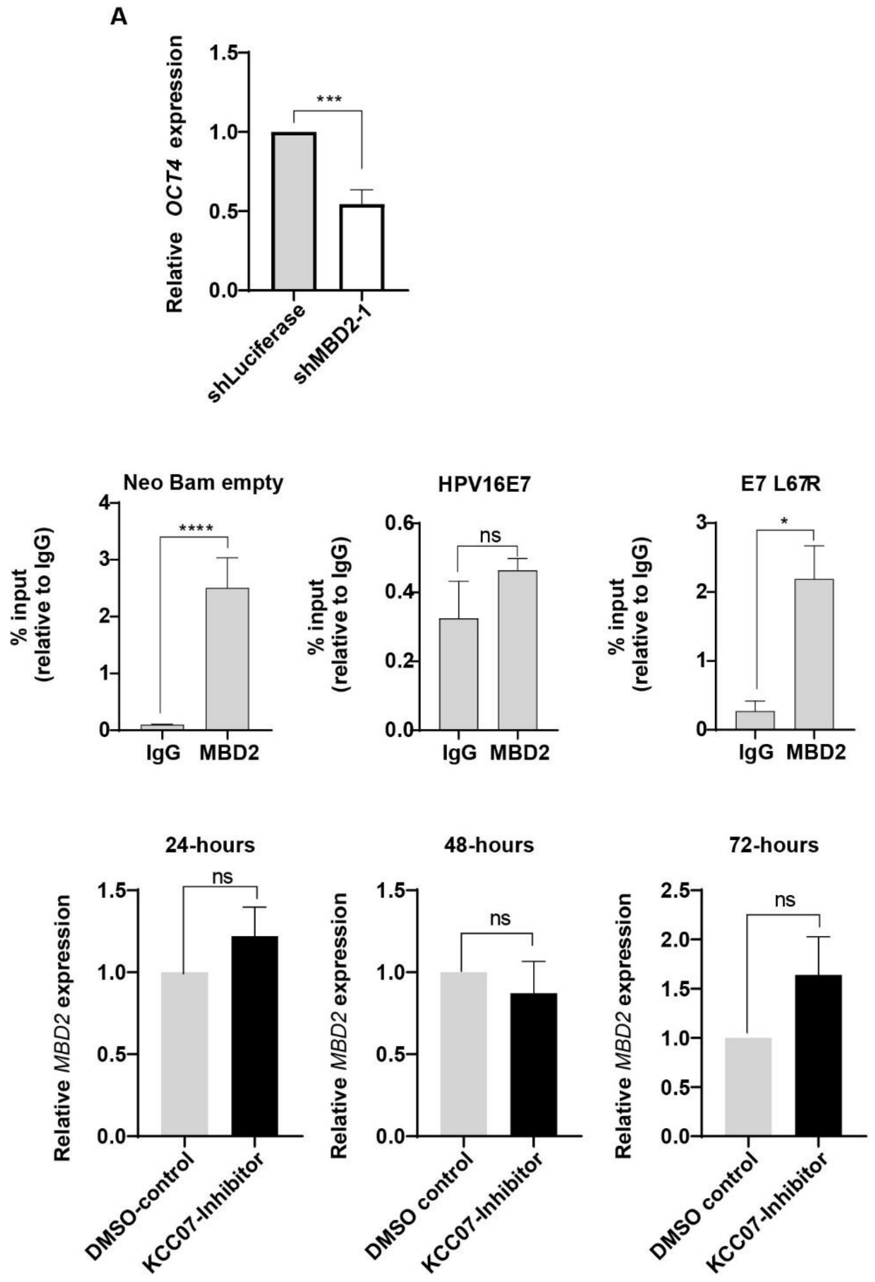


**Fig. S5.**

**KCC07 inhibition affects the binding of MBD2 on the hOct4 gene.** (A) MBD2-knockdown and the shLuciferase C33A cells were collected for RNA extraction to investigate the expression profile of Oct4. GAPDH was used as the housekeeping gene. (B) Chromatin immunoprecipitation was performed on C33A cells expressing the Neo Bam Empty, HPV16 E7 and E7 L67R vectors. MBD2 was immunoprecipitated with the MBD2 antibody and IgG was used as the control. qRT-PCR was performed by using primers for the hOct4 gene. (C) The KCC07 inhibitor was applied at 250nM concentration on C33A cells to check the MBD2 mRNA expression for 24-, 48, and 72-hours post-administration. ACTIN was used as the housekeeping control. Three independent replicates were used and plotted values on graphs are the Mean±SEM. The statistical analysis was calculated with two-tailed Unpaired T-test with $p<0.05$ (ns = non-significant, *$p<0.05$, **$p<0.01$, ***$p<0.001$, ****$p<0.0001$).

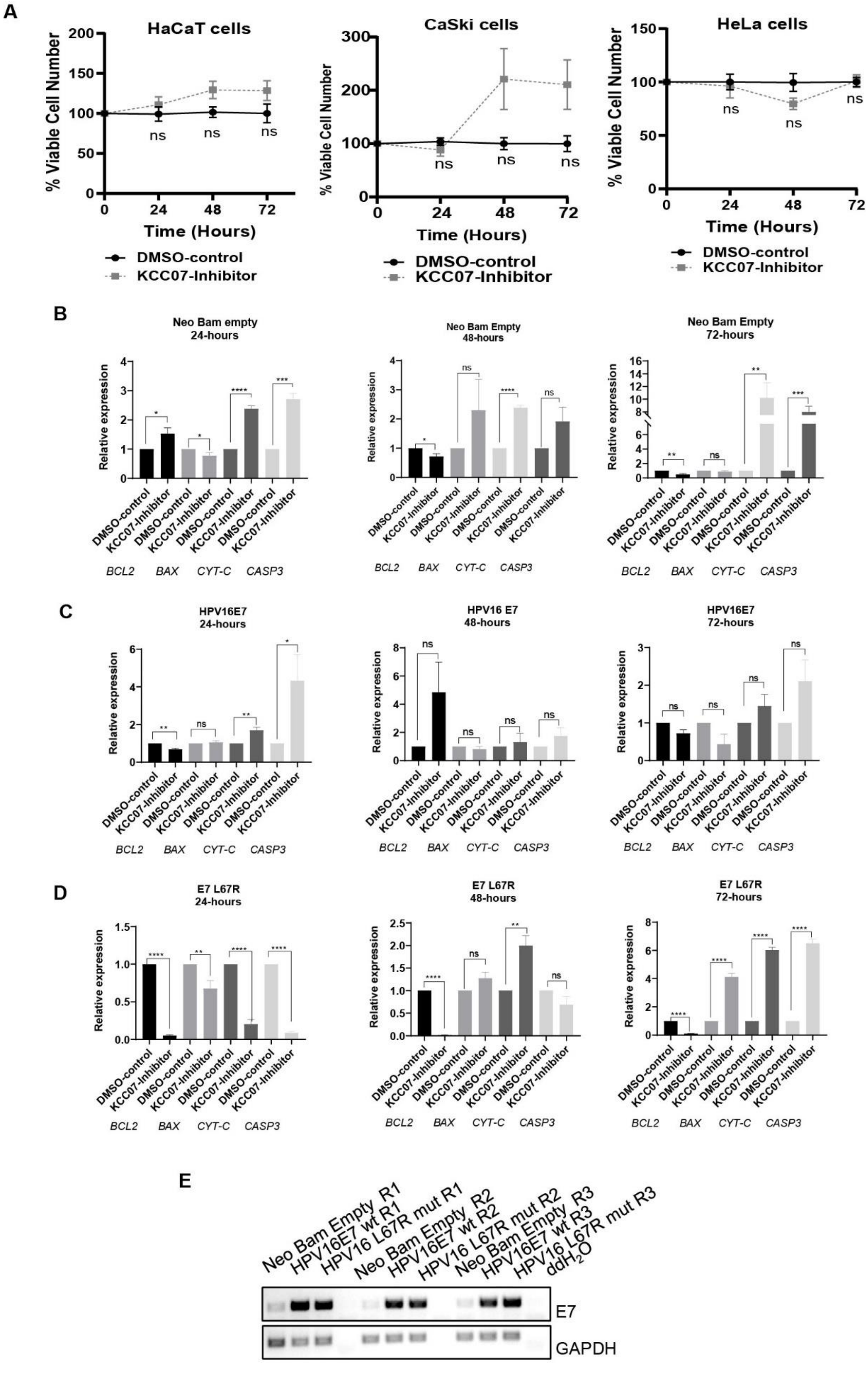
A
HaCaT cells
CaSki cells
HeLa cells
% Viable Cell Number
Time (Hours)
DMSO-control
KCC07-Inhibitor
B
Neo Bam empty 24-hours
Neo Bam Empty 48-hours
Neo Bam Empty 72-hours
Relative expression
BCL2 BAX CYT-C CASP3
C
HPV16E7 24-hours
HPV16 E7 48-hours
HPV16E7 72-hours
D
E7 L67R 24-hours
E7 L67R 48-hours
E7 L67R 72-hours
E
Neo Bam Empty R1
HPV16E7 wt R1
HPV16 L67R mut R1
Neo Bam Empty R2
HPV16E7 wt R2
HPV16 L67R mut R2
Neo Bam Empty R3
HPV16E7 wt R3
HPV16 L67R mut R3
ddH2O
E7
GAPDH

**Fig. S6.**

**MBD2 inhibition increases apoptosis in C33A cells not expressing HPV16 E7.** (A) HPV-positive cervical cancer cells (HeLa and CaSki) and human immortalized keratinocytes (HaCaT) were treated with the inhibitor at 250nM while DMSO was used as the negative control. Cell viability was assessed for 24-, 48- and 72-hours post-application of the inhibitor and was calculated by measuring cell numbers with a hemocytometer. C33A cells transfected with the (B) Neo Bam empty, (C) HPV16 E7 and (D) E7 L67R vectors and genes involved in the apoptotic pathway were investigated at the mRNA level at 24-, 48- and 72-hours post-administration of the inhibitor. Actin was used as the housekeeping gene. (E) RT-PCR was performed to validate the transfection of E7 plasmids in C33A cells and GAPDH was used as a loading control. Three independent replicates were used and plotted values on graphs are the Mean±SEM. The statistical analysis was calculated with two-tailed Unpaired T-test with $p<0.05$ (ns = non-significant, *$p<0.05$, **$p<0.01$, ***$p<0.001$, ****$p<0.0001$).

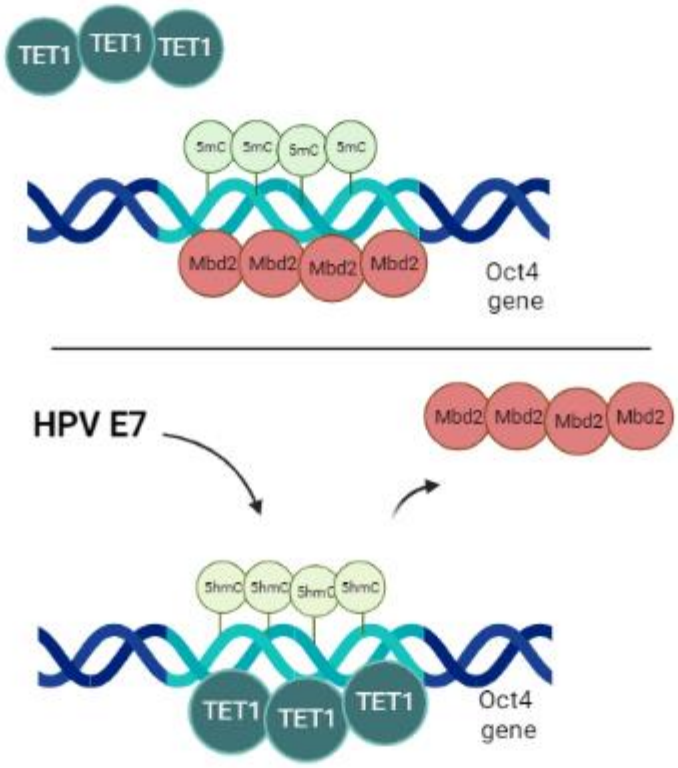


**Fig. S7.**

**E7 increases hydroxymethylation on the hOct4 gene via the displacement of MBD2.** Upon the absence of E7 in cervical cancer cells, MBD2 binds the hOct4 gene mediating an increase in methylation. When E7 is expressed in cervical cancer cells, MBD2 is displaced from hOct4 gene, TET1 expression increases and Oct4 becomes hydroxymethylated.

| | | |
|---|---|---|
| **Hs MBD2** | TTCAAGGAGTTGGTCCAGGTAG | GCAGGGTTCTTTTCCACAGC |
| **Hs MBD3** | CGGCCACAGGGATGTCTTTT | TGCTGGGGTGGTTGGTAATC |
| **Hs GAPDH** | TGCACCACCAACTGCTTAGC | GGCATGGACTGTGGTCATGAG |
| **Hs TET1** | AAGGGAGCCAACAAAAATGT | AGGACTCTGGGTTCTGAAAA |
| **Hs Sox2** | CGCCCCCAGGGGCAGCAGACTT CACA | CTCCTCTTTTGCACCCCTCCCATT T |
| **Hs KLF4** | GAAATTCGCCCGCTCAGATGATG AACT | TCTTCATGTGTAGTAACGAGGCGA GGTGGT |
| **Hs NANOG** | CCTGTGATTTGTGGGCCTGA | CTCTGCAGAAGTGGGTTGTTTG |
| **Hs C-MYC** | ATTCTCTGCTCTCCTCGACG | TGCGTAGTTGTGCTGATGTG |
| **Hs ALDH1A** | AACAACGAGTGGCAGAACTCAGAGAG | TTAGGGATTC CATGGTTGCA |
| **Hs N-CADHERIN** | GCGTCTGTAGAGGCTTCTGG | GCCACTTGCCACTTTTCCTG |
| **Hs SLUG** | TCCTCCATCTGACACCTCCTC | CCCCCGTGTGAGTTCTAATGT |
| **Hs FN** | GAGGAAACCTGCTCCAGTGC | CACGAACATCGGTGAAGGGG |
| **Hs CYT-C** | AAGGGAGGCAAGCACAAGACTG | CTCCATCAGTGTATCCTCTCCC |
| **Hs TP53** | CCTCAGCATCTTATCCGAGTGG | TGGATGGTGGTACAGTCAGAGC |
| **Hs CASP3** | GGAAGCGAATCAATGGACTCTGG | GCATCGACATCTGTACCAGACC |
| **Hs CASP9** | GTTTGAGGACCTTCGACCAGCT | CAACGTACCAGGAGCCACTCTT |
| **Hs BAX** | TCAGGATGCGTCCACCAAGAAG | TGTGTCCACGGCGGCAATCATC |
| **Hs BCL-2** | ATCGCCCTGTGGATGACTGAGT | GCCAGGAGAAATCAAACAGAGGC |
| **Hs THSB1** | TTGTCTTTGGAACCACACCA | CTGGACAGCTCATCACAGGA |
| **Hs RGS4** | ATG TGC AAA GGA CTC GCT GGT | TTA GGC ACA CTG AGG GAC TAG |
| **Hs ANKRD1** | AGACTCCTTCAGCCAACATGATG | CTCTCCATCTCTGAAATCCTCAGG |
| **Hs BZW2** | TTTCTGGACTCTACAGGCTCAA | ACCATCATCTATGCGCGTTCC |
| **Hs PTPRC** | CTTCAGTGGTCCCATTGTGGTG | CCACTTTGTTCTCGGCTTCCAG |
| **Hs ZNF483** | ACTGGAAAACCTCAGGAACCTA | CATGGCAATTCAACCCAC |
| **Hs CHRD** | CAGGAGTGGGGGACTAACC | CAGCACCTCAGCAAAGCCT |
| **Hs GLS2** | TGACTATAGTGGCGATGTCTCA | GTTCCATATCCATGGCTGACAA |
| **Hs TDG** | CATGCAGCAGTGAACCTTGTGG | GGTCATCCACTGCCCATTAGGA |
| **Hs DNMT3A** | CCTCTTCGTTGGAGGAATGTGC | GTTTCCGCACATGAGCACCTCA |
| **Hs DNMT3B** | CCATGAAGGTTGGCGACAA | TGGCATCAATCATCACTGGATT |
| **Hs GADD45A** | CGT TTT GCT GCG AGA ACG AC | GAA CCC ATT GAT CCA TGT AG |
| **Hs DNMT1** | CGGTTCTTCCTCCTGGAGAATGTCA | CACTGATAGCCCATGCGGACCA |
| **Hs PRMT6** | ACGAGTGCTACTCGGACGTT | AGTTCCGAAGGATACCCAGG |
| **Hs KDM1A** | TCAGGAGTTGGAAGCGAATCCC | GTTGAGAGAGGTGTGGCATTAGC |
| **Hs KDM2A** | CAAGGAGAGTGTGGTGTTTGCC | ACCTCTCCACAGAGGGAACATG |
| **Hs KDM2B** | CATGGAGTGCTCCATCTGCAATG | ACTTCGGACACTCCCAGCAGTT |
| **RT-PCR Primers** | | |
| **E7** | ATGGAGATACACCTACATTGCATGA | AATGGGCTCTGTCCGGTTCT |
| **Mycoplasma** | ACACCATGGGAGYTGGTAAT | CTTCWTCGACTTYCAGACCCAAGGCAT |

**Table S1.**

Primers used for qPCR and RT-PCR.

| Label | Sequence F (5'-3') | Sequence R (5'-3') |
|---|---|---|
| Seq 1 | TTGGATAGAATGTCCAAGCAGAGT | TGACAGAAAGGAGAATGACATTAGA |
| Seq 2 | CCATGTCTCTCTGCGGGC | AGTTGGAGGAGCCAGAGCTA |
| Seq 3 | AGCGAACCAGTATGGAGAACC | ATCTGCTGCAGTGTGGGTTT |
| Seq 4 | CCCCTCCGTCTTCCAGAATC | CCCCTCCGTCTTCCAGAATC |
| Seq 5 | CCAACCCCTTAGTCTGTTAGATGAG | CCCCACCCCTCCGTCTTC |
| Seq 6 | CTCTTCCCCCAGAAACTGGC | AGGCCAGGGTCTCTCTTTCT |
| Seq 7 | GTGGCTGAGGCCAGGG | CTTTCATGTCCTACATCCTATCCT |
| Seq 8 | CTTGGGGCGCCTTCCTTC | CATCACCTCCACCACCTGGA |
| Seq 9 | GAGTAGTCCCTTCGCAAGCC | GAGAAGGCGAAATCCGAAGC |
| Seq 10 | AGTAGTCCCTTCGCAAGCC | AAATCCGAAGCCAGGTGTCC |

**Table S2.**

Primers targeting different loci on the hOct4 gene.

| Vector | Catalog Number -Addgene | Type | Selection |
|---|---|---|---|
| pSMP-Luc | 36394 | Retroviral | Puromycin (2ug/ml) |
| pSMP-MBD2-1 | 36368 | Retroviral | Puromycin (2ug/ml) |
| pSMP-MBD2-2 | 36369 | Retroviral | Puromycin (2ug/ml) |
| pUMVC Packaging | 8449 | Retroviral | N/A |
| VSV-G Envelop | 14888 | Mammalian | N/A |
| pcDNA3.3_ OCT4 | 26816 | Mammalian | N/A |
| cmv- Neo Bam | 16440 | Mammalian | Neomycin (500ug/ml) |
| cmv 16 E7 | 13686 | Mammalian | Neomycin (500ug/ml) |
| cmv 16 E7 L67R | 13702 | Mammalian | Neomycin (500ug/ml) |

**Table S3.**

Plasmids used for this manuscript.

| Antibody | Company | Method for detection | Concentration |
|---|---|---|---|
| Oct4 | CST (2750S) | WB, IP | WB (1:800), IP (1:50) |
| E7 | Santa Cruz (ED17) | WB | WB (1:500) |
| TET1 | Abcam (ab272900) | WB | WB (1:800) |
| MBD2 | Abcam (ab188474) | WB, ChIP | WB (1:1000) ChIP (1:50) |
| MBD3 | CST (14540) | WB | WB (1:7500) |
| CHD4 | CST (11912) | WB | WB (1:500) |
| HDAC2 | CST (5113) | WB | WB (1:7500) |
| HDAC1 | CST (5356) | WB | WB (1:7500) |
| MTA1 | CST (5647) | WB | WB (1:7500) |
| RBBP7 | CST (6882) | WB | WB (1:7500) |
| 5hmC | Active Motif (39769) | DB, IF | DP (5:10000), IF (1:250) |
| 5Mc | Active Motif (61479) | IF | IF (1:250) |
| IgG | Merck (SKU 12-370) | IP, ChIP | IP (1:50) ChIP (1:50) |
| MCM7 | Abcam (ab2360) | WB | WB (1:1000) |
| Vimentin | Abcam (ab137321) | WB | WB (1:1000) |
| p53 | Abcam (ab26) | WB | WB (1:1000) |
| PCNA | Santa Cruz (sc-25280) | WB | WB (1:1000) |
| Gapdh | Abcam (ab9484) | WB | WB (1:1000) |
| **Secondary antibodies** | | | |
| mouse anti-rabbit IgG HRP | Santa Cruz (sc-2357) | WB | WB (1:1000) |
| mouse anti-goat IgG HRP | Santa Cruz (sc2354) | WB | WB (1:1000) |
| m-IgGk BP-HRP | Santa Cruz (sc-516102) | WB | WB (1:1000) |
| FITC anti-rabbit | Jackson ImmunoResearch | IF | IF (1:250) |

### Table S4.

List of antibodies used in this study. WB=western blot, IP=immunoprecipitation, ChIP=Chromatin Immunoprecipitation, IF=immunofluorescence, DB=Dot blot
